# Local adaptation to climate facilitates a global invasion

**DOI:** 10.1101/2024.09.12.612725

**Authors:** Diana Gamba, Megan L. Vahsen, Toby M. Maxwell, Nikki Pirtel, Seth Romero, Justin J. Van Ee, Amanda Penn, Aayudh Das, Rotem Ben-Zeev, Owen Baughman, C. Sean Blaney, Randy Bodkins, Shanta Budha-Magar, Stella M. Copeland, Shannon L. Davis-Foust, Alvin Diamond, Ryan C. Donnelly, Peter W. Dunwiddie, David J. Ensing, Thomas A. Everest, Holly Hoitink, Martin C. Holdrege, Ruth A. Hufbauer, Sigitas Juzėnas, Jesse M. Kalwij, Ekaterina Kashirina, Sangtae Kim, Marcin Klisz, Alina Klyueva, Michel Langeveld, Samuel Lutfy, Daniel Martin, Christopher L. Merkord, John W. Morgan, Dávid U. Nagy, Jacqueline P. Ott, Radoslaw Puchalka, Lysandra A. Pyle, Leonid Rasran, Brian G. Rector, Christoph Rosche, Marina Sadykova, Robert K. Shriver, Alexandr Stanislavschi, Brian M. Starzomski, Rachel L. Stone, Kathryn G. Turner, Alexandra K. Urza, Acer VanWallendael, Carl-Adam Wegenschimmel, Justin Zweck, Cynthia S. Brown, Elizabeth A. Leger, Dana M. Blumenthal, Matthew J. Germino, Lauren M. Porensky, Mevin B. Hooten, Peter B. Adler, Jesse R. Lasky

## Abstract

Local adaptation may facilitate range expansion during invasions, but the mechanisms underlying successful invasions remain unclear. Cheatgrass (*Bromus tectorum*), native to Eurasia and Africa, has invaded globally, with severe impacts in western North America. We aimed to identify mechanisms and consequences of local adaptation in the North American cheatgrass invasion. We sequenced 307 range-wide genotypes and conducted controlled experiments. We found that diverse lineages invaded North America, where long-distance gene flow is common. Nearly half of North American cheatgrass comprises a mosaic of ∼19 locally adapted, near-clonal genotypes, each seemingly very successful in a different part of North America. Additionally, ancestry, phenotype, and allele frequency-environment clines in the native range predicted those in the invaded range, indicating pre-adapted genotypes colonized different regions. Common gardens showed directional selection on flowering time that reversed between warm and cold sites, potentially maintaining clines. In the USA Great Basin, genomic predictions of strong local adaptation identified sites where cheatgrass is most dominant. Our results indicate that multiple introductions and migration within the invaded range fueled local adaptation and success of cheatgrass in western North America. Understanding how environment and gene flow shape adaptation and invasion is critical for managing ongoing invasions.

## INTRODUCTION

Biological invasions are a major cause of global biodiversity decline and ecosystem disruption, but the mechanisms driving ongoing invasions remain poorly understood^1,2^. In particular, the role of adaptive evolution in enabling invasive species to succeed is poorly understood. Is invasion success primarily determined by the susceptibility of invaded ecosystems^3^, or are the worst invaders adapted to spread and dominate^4^? For example, local adaptation to invaded environments could increase fitness and abundance^5^, while ongoing gene flow into the invaded range could swamp or reshape local adaptation^6,7^. If colonizing propagules are diverse, new populations may quickly adapt to local environments, facilitating invasive spread^4,8^. However, colonizing genotypes may reach new environments to which they are maladapted, swamping local adaptation and potentially hindering further spread^6,7,9,10^. Furthermore, if the diversity of colonizing propagules is low, new populations may be unable to adapt to new conditions^11^, phenotypic plasticity of invasive genotypes may counteract local adaptation^1^, and/or colonization bottlenecks may increase the frequency of deleterious mutations^12^. Testing these hypotheses requires a rare combination of genomic, fitness, and abundance data^13^.

Multiple mechanisms could contribute to local adaptation during invasions, generating distinct patterns of genomic and phenotypic variation^14,15^. In general, selection may change along environmental gradients and promote genotypic and phenotypic clines^16^. If environmental gradients are similar in native versus invaded regions, clines may be similar between native and invaded regions, indicating niche conservatism of different lineages^17–19^. Alternatively, if selective pressures are novel in the invaded range, clines may be distinct between native and invaded regions, suggesting niche shift of some lineages^19–22^. Furthermore, invasive genotypes may closely match the genetic diversity of the native range^23^, represent newly admixed populations^24^, or form novel genotypes via introgression from congeners^25^. Understanding successful invasions thus requires dissecting global patterns of genomic and phenotypic variation, which has seldom been accomplished. Although some studies have examined genomic and phenotypic differences between native and invasive populations, sampled populations between ranges are hard to compare because they often differ in spatial scale and/or do not incorporate enough environmental variation^26^.

*Bromus tectorum* L. (cheatgrass) is a grass native to Eurasia and northern Africa that spread across North America by the 1890s^27,28^, heavily influencing ecological dynamics of arid and semi-arid ecosystems of the North American Intermountain West^29^. At least some introductions likely came via contamination in grain shipments^27,28^. Cheatgrass occurs in high abundance across an estimated 31% (210,000 km^2^) of this region^30^, displacing native perennials via rapid reproduction and shortened fire return intervals^29^, reducing biodiversity and degrading wildlife habitat^31^. It is highly selfing, typically winter annual, with a high-quality reference genome (∼2.5 Gb)^28,32^. Existing genetic studies, while limited to small numbers of markers or populations, suggest that multiple introductions from different regions in Europe might have occurred in North America^28,33–38^. Studies have shown evidence for local adaptation in phenology at small scales^39–43^, and substantial genetic differentiation in phenology between populations from different regions^32,44^, although range-wide patterns of local adaptation remain elusive. Due to cheatgrass’s high rate of selfing, novel recombinant genotypes are expected to be rare, limiting novel genomic diversity in the invaded range. However, repeated introductions into North America and post-introduction dispersal could have promoted the adaptative potential and invasive spread of cheatgrass populations.

Here, we aim to identify mechanisms and consequences of local adaptation in the North American cheatgrass invasion. We hypothesize that for cheatgrass, as a self-pollinating species with few constraints on dispersal, local adaptation in the invaded range would be likely due to rapid spread of pre-adapted genotypes to suitable environments (as opposed to adaptation by *de novo* mutation or novel admixtures), provided there were multiple diverse introductions. We sequence whole genomes of a global panel of 307 genotypes from the native and invaded ranges and measure phenotypes and performance in one growth chamber experiment and two field common gardens. We ask whether there were multiple and diverse introductions to North America and examine genetic consequences of the invasion. We evaluate how geography, environment, and phenotype shape genomic diversity. We test whether ancestry, trait, and allele frequency-environment clines were repeated in native and invasive genotypes, and if selection maintains clines. Finally, we integrate field surveys of cheatgrass abundance in the USA Great Basin^30^ to assess whether genomic matching to local climates facilitated invasive dominance. Our results reveal that multiple introductions and migration within the invaded range fueled local adaptation and success of cheatgrass in western North America.

## RESULTS AND DISCUSSION

### Diverse native range ancestries invaded North America

Cheatgrass populations in North America stem from multiple, diverse introductions. Using ∼267k unlinked single-nucleotide polymorphisms (SNPs), different clustering analyses of global genomic variation showed that population genetic structure largely followed geography in the native range and to a lesser degree in North America (Fig. 1 showing K=4 ancestral genetic clusters/ancestries, Supplementary Fig. 1). In the native range, west Asian, Mediterranean, and Atlantic genotypes primarily fell in a single ancestry, while central and eastern European genotypes were mostly assigned to two ancestries differentiated by latitude and were overall more intermediate (*i.e.*, composed of multiple ancestries). In the invaded range, genotypes were assigned to all four ancestries in western North America (WNA, west of the Rocky Mountains), but only to two ancestries in eastern North America (ENA, east of the Rocky Mountains) (Fig. 1a–c, Supplementary Fig. 2). The majority of invasive genotypes were similar to genotypes from north, central, or eastern Europe (Fig. 1d–f). In WNA, however, warm desert genotypes in southern California and Nevada were similar to genotypes from Iran and Afghanistan. The warm Mojave and the cool Pacific Northwest also harbored genotypes similar to those from the western Mediterranean (Fig. 1e,f). For regions with less extensive invasions, results showed that: Argentines are similar to Spanish genotypes, an Australian is similar to western Mediterranean genotypes and a widespread lineage from WNA, New Zealand genotypes are similar to northeastern European genotypes, and a Korean genotype is similar to central eastern European genotypes and a widespread lineage from ENA.

**Fig. 1:**
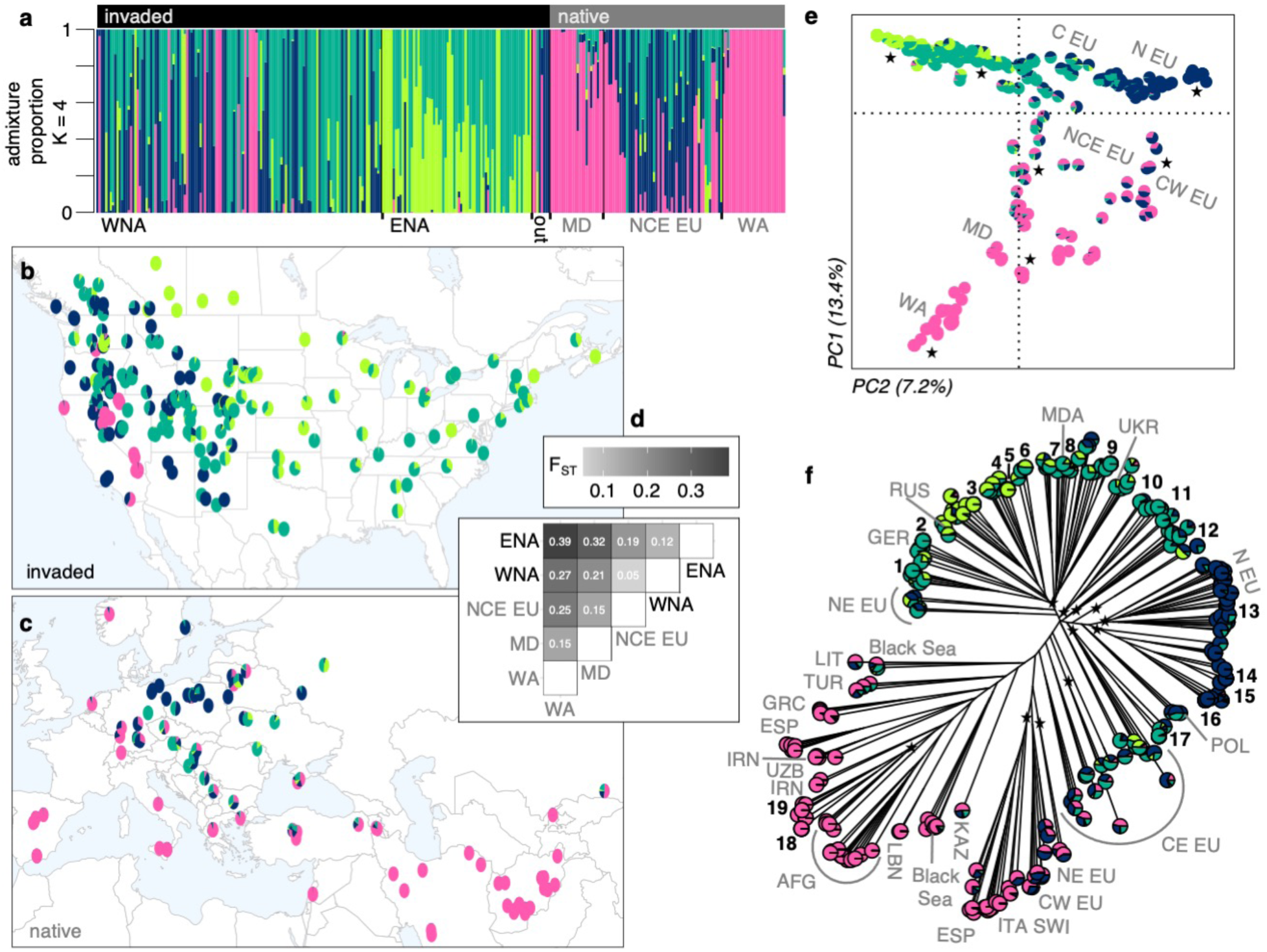
The cheatgrass invasion involved multiple diverse introductions from the native range to North America. (**a**) Admixture proportions for K=4 ancestral genetic clusters (colors) for invasive and native genotypes in different regions; WNA: western North America (n=107), ENA: eastern North America (n=67), out: not in North America (n=8), MD: Mediterranean (n=24), NCE EU: north-central-east Europe (n=53), WA: west Asia (n=28). Geographic distribution of (**b**) invasive (n=194, North American only) and (**c**) native (n=105) genotypes. (**d**) Genetic differentiation (F_ST_) between native and invaded regions, with notations following panel a. (**e**) Principal components analysis showing PC1 (y-axis) and PC2 (x-axis) explaining 20.6% of genomic variation. Axes are shifted to better reflect the latitudinal distribution of genotypes. Gray letters denote geographic origin in the native range and stars represent genotypes in the invaded range. (**f**) Neighbor-joining tree annotated with native (gray letters) and invaded locations (black numbers and stars). Native notations follow the ISO alpha-3 country code or their cardinal direction in Europe (EU). Black numbers mark groups of 2–14 near-clonal, and often widely distributed, invasive genotypes. Stars mark branches with invasive genotypes.

The diversity of genotypes found in WNA reflects colonization by propagules from different native regions, while patterns in ENA reflect reduced genetic diversity. Accordingly, population-specific F_ST_ (*i.e.*, the degree of relatedness among individuals) was higher in the invaded compared to the native range (0.2 and 0.03, respectively) especially for ENA (0.39) compared to WNA (0.18). Pairwise F_ST_ values were lowest between European and North American genotypes, while other pairs of regions were more diverged (e.g., those involving the Mediterranean and west Asia, Fig. 1d). The native and invaded range were moderately genetically differentiated (F_ST_ = 0.11), comparable to the differentiation between genotypes from ENA and WNA (F_ST_ = 0.12). In the native range, pairwise F_ST_ values were larger (Fig. 1d), showing strong divergence between European and west Asian genotypes (F_ST_ = 0.25).

### WNA and ENA: different patterns of diversity

Much of North America harbors great genomic diversity with little evidence of elevated genetic load and inbreeding compared to the native range (Supplementary Figs. 3–5 using ∼15.1M SNPs). In WNA, nucleotide diversity (π, Supplementary Fig. 3a,b) was comparable to the most diverse native region, north-central-eastern Europe (0.0016 ± 4.5×10^−6^ se vs. 0.0018 ± 4.6×10^−6^ se, respectively), followed by the Mediterranean (0.0015 ± 3.8×10^−6^ se) and west Asia (0.0011 ± 3.1×10^−6^ se). Nucleotide diversity was much lower in ENA (0.0009 ± 4.5×10^−6^ se). In WNA, the skew in the site frequency spectrum (Tajima’s D, Supplementary Fig. 3c,d) was positively shifted (mean=2.8 ± 0.006 se), indicating an excess of intermediate-frequency SNPs, consistent with strong population structure and heterogeneous ancestry across the region (see also^38^). In ENA, Tajima’s D was low (mean=0.5 ± 0.009 se), indicating more rare variants and suggesting recent population expansion. In the Mediterranean and north-central-eastern Europe, Tajima’s D was positively shifted (Mediterranean mean=1.6 ± 0.005, north-central-eastern Europe mean=2.2 ± 0.007), reflecting substantial population structure within these regions. The Mediterranean comprises multiple distinct eastern and western lineages (see also^45^), while north-central-eastern Europe comprises multiple lineages with some intermediate genotypes. In contrast, west Asian genotypes appeared more closely related to each other (Tajima’s D mean=0.3 ± 0.006).

To understand the effects of potential bottlenecks and drift in North America, we first examined deleterious mutation load using ∼15.1M SNPs, under the hypothesis that most protein changing mutations are deleterious. Estimated mutation load was not different between native versus invaded range genotypes of the same ancestry (two-way ANOVA: range F_(1,290)_=57.8, *p*=4×10^−13^, ancestry F_(4,290)_=46.7, *p*<2×10^−16^, interaction F_(3,290)_=4.4, *p*=0.005; Tukey HSD range *p*=0.2, Supplementary Fig. 4). The central-eastern European ancestry (teal in Supplementary Fig. 4), widespread in North America, showed the lowest load in both ranges, suggesting large effective population size at some point in the past. In contrast, the west Asian and Mediterranean ancestry (pink in Supplementary Fig. 4) was associated with higher load in both ranges.

Next we examined runs of homozygosity (ROH^46^) using a panel of 101 closely related native and invasive genotypes sequenced directly from field collections (Supplementary Fig. 5). Native and invasive genotypes (grouped by range) had similar Tajima’s D, thus similar skew in the site frequency spectrum. The native group, however, had lower counts of ROH and a much higher FROH (the proportion of the genome with ROH), resulting in inference of a strong selfing rate (Supplementary Fig. 5a–c). Selfing rates were significantly different between the native and invasive groups (two-tailed t-test t=3.8437, df=11.896, *p*=0.002), though both were >0.9 (Supplementary Fig. 5c). This could reflect relaxed selection for reproductive assurance in invasive genotypes from specific environments. For example, although selfing is more common, some highly inbred desert lineages appeared overrepresented among parents of heterozygotes in a previous common garden experiment^47^. Our results suggest that the North American cheatgrass invasion is not associated with higher inbreeding due to selfing compared to the native range (Supplementary Fig. 5c,d).

Taken together, the high diversity in WNA indicates great potential for adaptation in this heavily invaded region. In contrast, the lower diversity in ENA reflects colonization by a few closely related lineages (see also^34^) that persist as ruderal plants in urban and agricultural environments.

### Strong isolation-by-environment in North America

Both geography and environment shape genomic diversity in the native range, but geography plays a weak role in North America. Isolation-by-distance (based on ∼267k SNPs) was strong in the native range (geographic vs. genetic distance Mantel *p*=10^−4^, Fig. 2a) but very weak in North America (Mantel *p*=0.06, Fig. 2b). At 0–100 km distance, pairs of distantly related genotypes were common in WNA, but not in ENA or the native range (Supplementary Fig. 6a,b). Even at the smallest scales (0–25 km), isolation-by-distance appeared weaker in the invaded compared to the native range (Supplementary Fig. 6c,d). Moreover, several groups in North America (“1–19” in Fig. 1f) composed of 2–14 near-clonal genotypes (>98% SNPs identity) were found across distances of >3000 km (Fig. 2b, Supplementary Fig. 6a, 7a). In contrast, such widely distributed, near-clonal genotypes were absent in the native range (Fig. 2a, Supplementary Fig. 6b). These patterns suggest long-distance dispersal within North America by lineages descended from distinct native range populations. Furthermore, groups of near-clonal genotypes occupied significantly different environments (PERMANOVA of multivariate environment predicted by clonal group: *p*=0.0001, R^2^=0.65) that together encompass the extent of climate space in North America (Supplementary Fig. 7b). This suggests that although genotypes might not be dispersal limited in North America, their spread may be limited by different environmental constraints, suggesting local adaptation. The weak spatial patterns in North America may also reflect genotype sorting along the steep, heterogeneous climatic gradients that are common in WNA. Pairwise climatic distance (based on Euclidian distance in Supplementary Fig. 6e) significantly increased with spatial distance in both native and invaded ranges (Mantel *p*=10^−4^ and Mantel Pearson correlation=0.6 in both ranges; Supplementary Fig. 6f,g), but this relationship was weak in WNA (Mantel Pearson correlation=0.3 WNA vs. 0.7 ENA), reflecting the fine-scale climatic heterogeneity of this region.

**Fig. 2:**
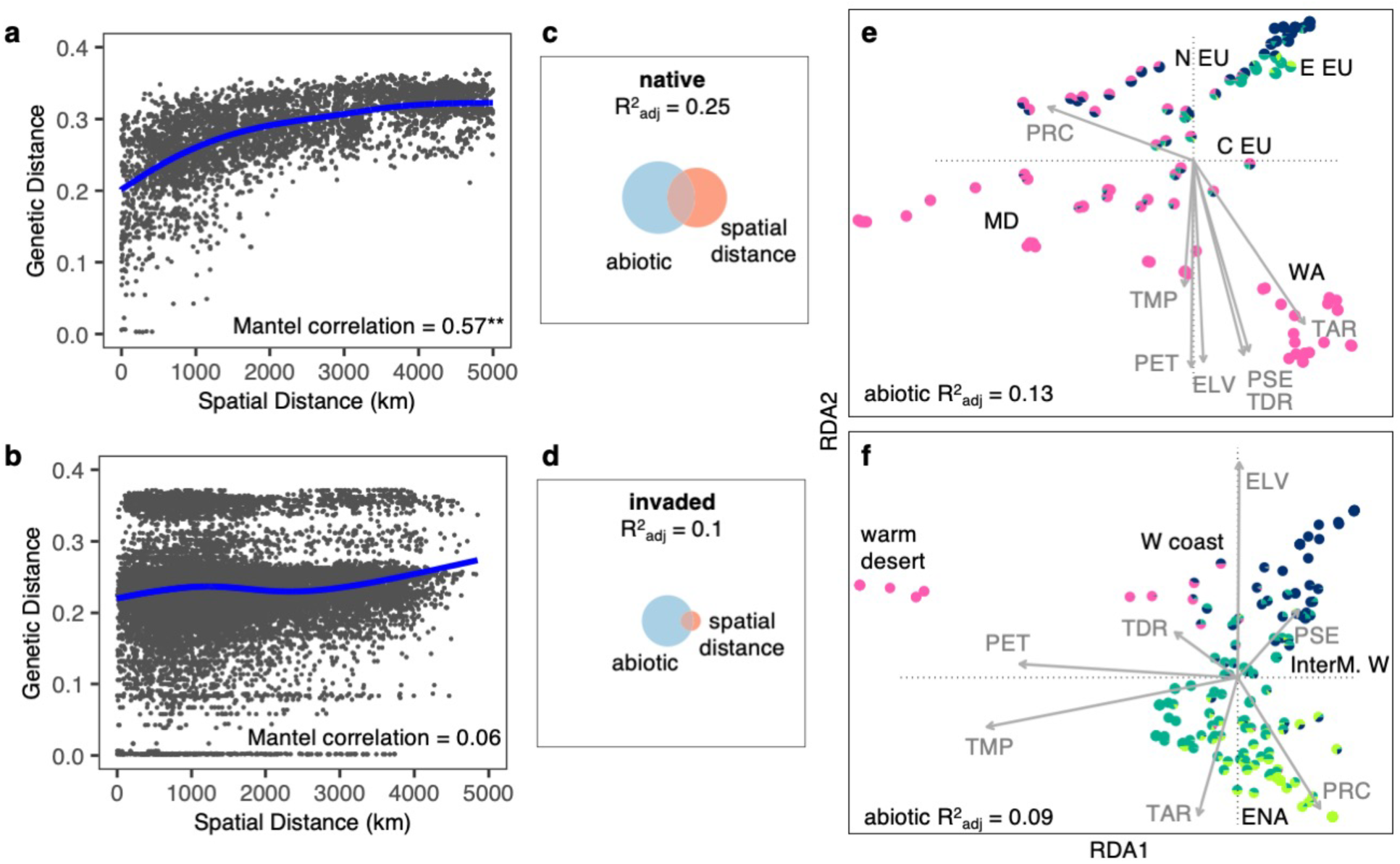
Genomic variation is structured by environment in the native and invaded ranges. Strong isolation-by-distance in the (**a**) native (**Mantel *p*=10^-4^) but not in the (**b**) invaded range; plots show raw pair-wise data with a spline. Euler Plots show genomic variation is best explained by both the abiotic environment and spatial distance in (**c**) the native range, but only by the abiotic environment in (**d**) the invaded range. Fields of squares represent total genomic variation, circles represent genomic variation explained by a particular group of variables calculated using variance partitioning with RDA ordination (native n=105, invaded n=194). (**e**) Native and (**f**) invasive genotypes projected on the first two canonical axes of RDA (x-axis: RDA1 y-axis: RDA2). Arrows represent environmental predictors that strongly correlate with a maximal proportion of variation in linear combinations of SNPs. ELV: elevation, PET: potential evapotranspiration, PRC: total annual precipitation, PSE: precipitation seasonality, TAR: temperature annual range, TDR: temperature diurnal range, TMP: annual mean temperature. Colors are K=4 ancestral clusters. Geographic annotations are depicted in bolded black; N EU: north Europe, E EU: east Europe, C EU: central Europe, MD: Mediterranean, WA: west Asia, W coast: west coast, InterM. W: intermountain west, ENA: eastern North America.

To further examine genomic differentiation along climate gradients, we performed redundancy analysis (RDA) with variance partitioning, comparing the role of climate and spatial variables in explaining genomic variation. SNP variation was better explained by these predictors in the native than in the invaded range (native R^2^_adj_=0.25, invaded R^2^_adj_=0.10; Fig. 2c,d). Spatial variables explained little SNP variation in North America (native R^2^_adj_=0.07, invaded R^2^_adj_=0.005), confirming low isolation-by-distance. In both ranges the abiotic environment explained the largest portion of SNP variation (native R^2^_adj_=0.13, invaded R^2^_adj_=0.09; Fig. 2e,f), highlighting the importance of isolation-by-environment in both the native and invaded range and consistent with invasive local adaptation via native pre-adaptation^19^.

### Repeated ancestry-climate clines

Ancestry-environment clines were remarkably similar in the native and invaded ranges, suggesting environmental filtering of pre-adapted genotypes that could disperse long distances or via directed gene flow (as opposed to local adaptation by novel genotypes). We focused on aridity and temperature gradients representative of global climatic variation in the cheatgrass range (see Supplementary Fig. 6e) and used generalized-additive-models (GAMs) to detect significant ancestry-climate trends between ranges (Supplementary Fig. 8). In native and invasive genotypes, the west Asian and Mediterranean genetic cluster (pink) was more frequent in drier regions (GAM *p*=0.0004, pseudo-R^2^=0.5), the northern Europe cluster (blue) was more frequent in humid regions (GAM *p*=0.007, pseudo-R^2^=0.08), the central Europe cluster (teal) was more frequent in regions with little precipitation seasonality (GAM *p*=10^−5^, pseudo-R^2^=0.2), and the presumably northeast Europe ancestry (green) was more frequent in regions with colder winters (GAM *p*=0.002, pseudo-R^2^=0.1).

### Repeated phenotype-climate clines

Consistent with the hypothesis that pre-adaptation to local climate facilitated the cheatgrass invasion, we found similar phenotype-environment clines in the invaded and native ranges. A principal components (PC) analysis on genetic variation among 169 native and invasive genotypes for eleven growth chamber phenotypes (Supplementary Data 1, Supplementary Table 1) detected multi-trait axes of variation (Fig. 3a). PC1 explained 35.4% of the variation and suggested a life history axis of delayed flowering and high vegetative investment (more tillers and leaves) versus rapid flowering and high reproductive investment (taller, more fecund inflorescences). PC2 explained 22.2% of the variation and indicated an axis associated with larger plants with greater growth after vernalization versus shorter plants with little growth after vernalization. Native genotypes had on average earlier flowering (two-tailed t-test t=4.09, df=41.36, *p*=0.0002) and higher reproductive investment (two-tailed t-test t= –1.92, df=43.69, *p*=0.06) than invasive genotypes (PC1 two-tailed t-test t=3.02, df=38.14, *p*=0.004) which may be due to different ancestry proportions in the native range. We found no significant native versus invasive trait differences after accounting for relatedness (*p*>0.4), thus no evidence for evolution of increased competitive ability by invasive cheatgrass^48^. Additionally, near-clonal groups were significantly different in multivariate phenotypes (PERMANOVA of multivariate phenotype predicted by clonal group: *p*=0.03, R^2^=0.39), yet the remaining non-clonal genotypes were also diverse (Supplementary Fig. 7c). These results highlight how North America hosts diverse life histories.

**Fig. 3:**
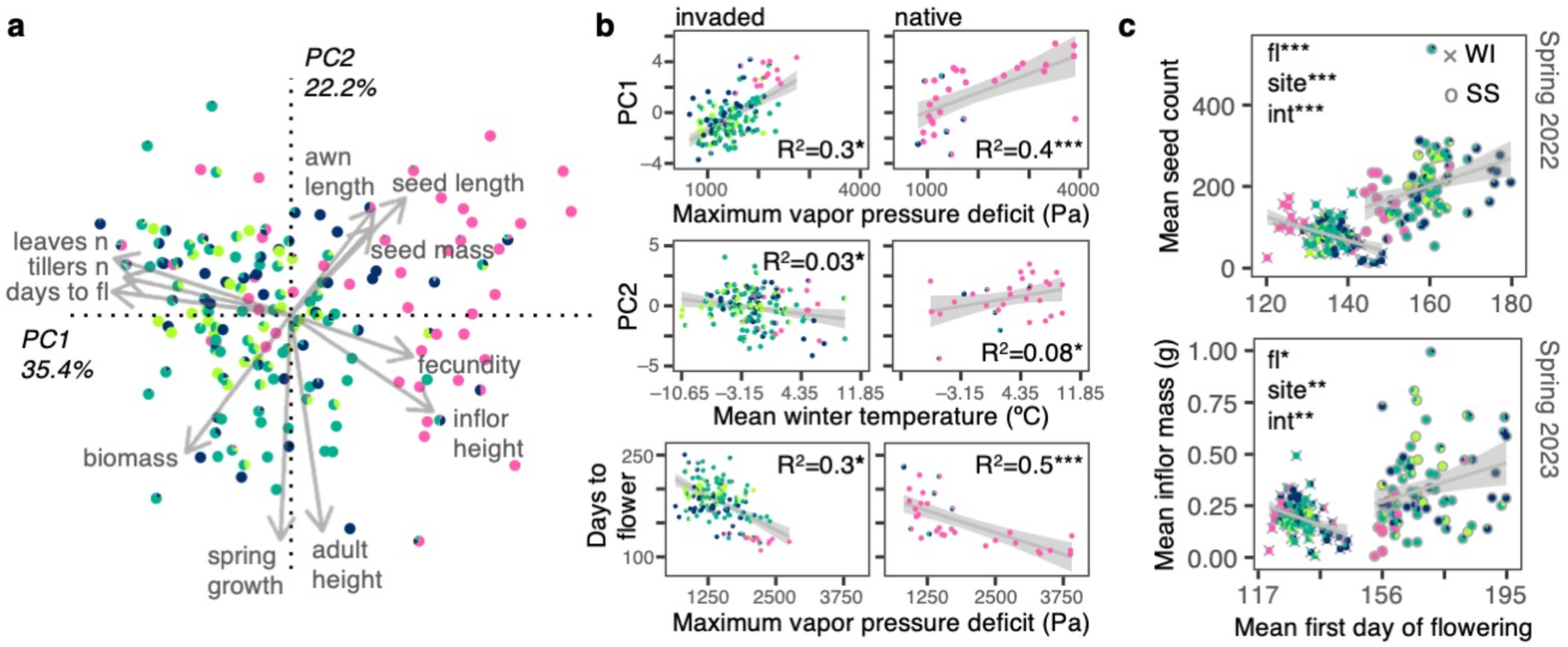
Selection along aridity and temperature gradients shapes flowering phenology. (**a**) Eigenvector plot with loadings of eleven phenotypes onto PC1 (x-axis) and PC2 (y-axis) describing axes of life history variation of 169 genotypes in a growth chamber; fl: Flowering, n: Number, inflor: Inflorescence. (**b**) Growth chamber phenotype-environment associations for invasive (left; n=138–145) and native genotypes (right; n=31–36). Coefficients of determination (R^2^), trends (gray lines), and 95% confidence intervals (gray shades) come from linear regressions. Significance comes from linear-mixed kinship models that accounted for relatedness among genotypes: \**p*<0.05, \*\*\**p*<0.0001. (**c**) Fitness advantage of early flowering genotypes at a warm site/common garden (WI: Wild Cat, gray crosses, n=93) and of late flowering genotypes at a cool site/common garden (SS: Sheep Station, gray open circles, n=82) in two consecutive years (top: 2022 Spring harvest and bottom: 2023 Spring harvest). Trends (gray lines) and 95% confidence intervals (gray shades) come from linear regressions. Significance comes from linear-mixed kinship models of fitness (seed count for 2022 and inflorescence mass for 2023) in response to mean first day of flowering (fl), site, and their interaction (int): \**p*=0.006, \*\**p*<0.0006, ***p<0.00001. In all panels colors are K=4 ancestral clusters.

Multiple trait-climate clines potentially maintained by selection were mirrored between the native and invaded range (Fig. 3b), indicating sorting of genotypes along humidity and temperature gradients. We focused on two climate variables that we hypothesized would capture distinct climatic stressors: maximum vapor pressure deficit (Pa), for drought adaptation, and mean winter temperature (°C), for cold adaptation (Supplementary Fig. 6e). To test for evidence of selection maintaining clines, we used linear-mixed models that accounted for genomic similarity (lmkin below), similar to Q_ST_–F_ST_ tests^49^. When significant, these models suggest selection is driving trait-climate clines, because the cline is stronger than expected by the genome-wide patterns of variation. In native and invasive genotypes, earlier flowering was associated with higher aridity (native lmkin-*p*=2e^−8^, R^2^=0.5; invaded lmkin-*p*=0.03, R^2^=0.3), suggesting a locally adaptive cline of rapid phenology/early reproductive investment in arid regions versus delayed phenology/early vegetative investment in humid regions. Also, clines showed evidence of selection specifically within WNA, but not in ENA (Supplementary Fig. 9). These patterns suggest local adaptation via life history clines in WNA. In contrast, life history in ENA is not associated with climate, potentially because a single generalist ruderal strategy is adaptive throughout ENA.

### Selection on flowering time along a temperature gradient in WNA

In field common gardens, selection on flowering time changed direction between sites that differed in temperature. To test whether phenotypic clines in WNA were promoted by selection, we conducted two common garden experiments in different climates in Idaho with fall plantings across two years (2021 and 2022). One site was cooler (Sheep Station, ID, USA 44.2456°N, 112.2144°W, annual mean temperature 6°C) and the other warmer (Wildcat, ID, USA 43.4744°N, 116.9018°W, annual mean temperature 12°C). We planted 95 diverse genotypes from across WNA, for a total of 14,800 plants^50^. We measured flowering time, survival, and fecundity. In both years (Fig. 3c), selection favored later flowering at the cool site, with late flowering genotypes having ∼3× the fitness of earlier flowering genotypes (∼300 vs. ∼100 seeds produced per original sown seed). By contrast, at the warm site, selection favored earlier flowering, with the earliest flowering genotypes having ∼10× the fitness of the later flowering genotypes (∼170 vs. ∼17 seeds produced per original sown seed). This suggests that late flowering genotypes have an extreme disadvantage in warm climates. This strong selection is consistent with our finding that the hottest sites in WNA were almost exclusively comprised of west Asian-like genotypes. We saw no clear admixture from distantly related, but geographically proximate European-like genotypes inhabiting cooler and wetter higher elevations in WNA (Fig. 1a), suggesting a barrier to dispersal of maladapted genotypes. Thus, climate gradients in WNA appear to impose changes in selection maintaining a strong phenotypic cline.

### Repeated allele frequency-climate clines

Putative quantitative trait loci (QTL) for traits under selection showed similar allele frequency-climate clines in the invaded and native ranges (Fig. 4, Supplementary Figs. 10–11). Using genome-wide association studies (GWAS), we identified several genetic loci associated with variation in flowering time and number of tillers (Supplementary Data 2). We implemented two methods that accounted for population genetic structure: univariate mixed model LMM and multilocus mixed model MLMM. Of the 400 top GWAS SNPs (100 top SNPs × 2 phenotypes × 2 GWAS methods), just one SNP was segregating exclusively within the invaded range. The other 399 SNPs segregated in both native and invaded ranges, supporting the hypothesis that *de novo* mutations have not been major drivers of local adaptation in North America.

**Fig. 4:**
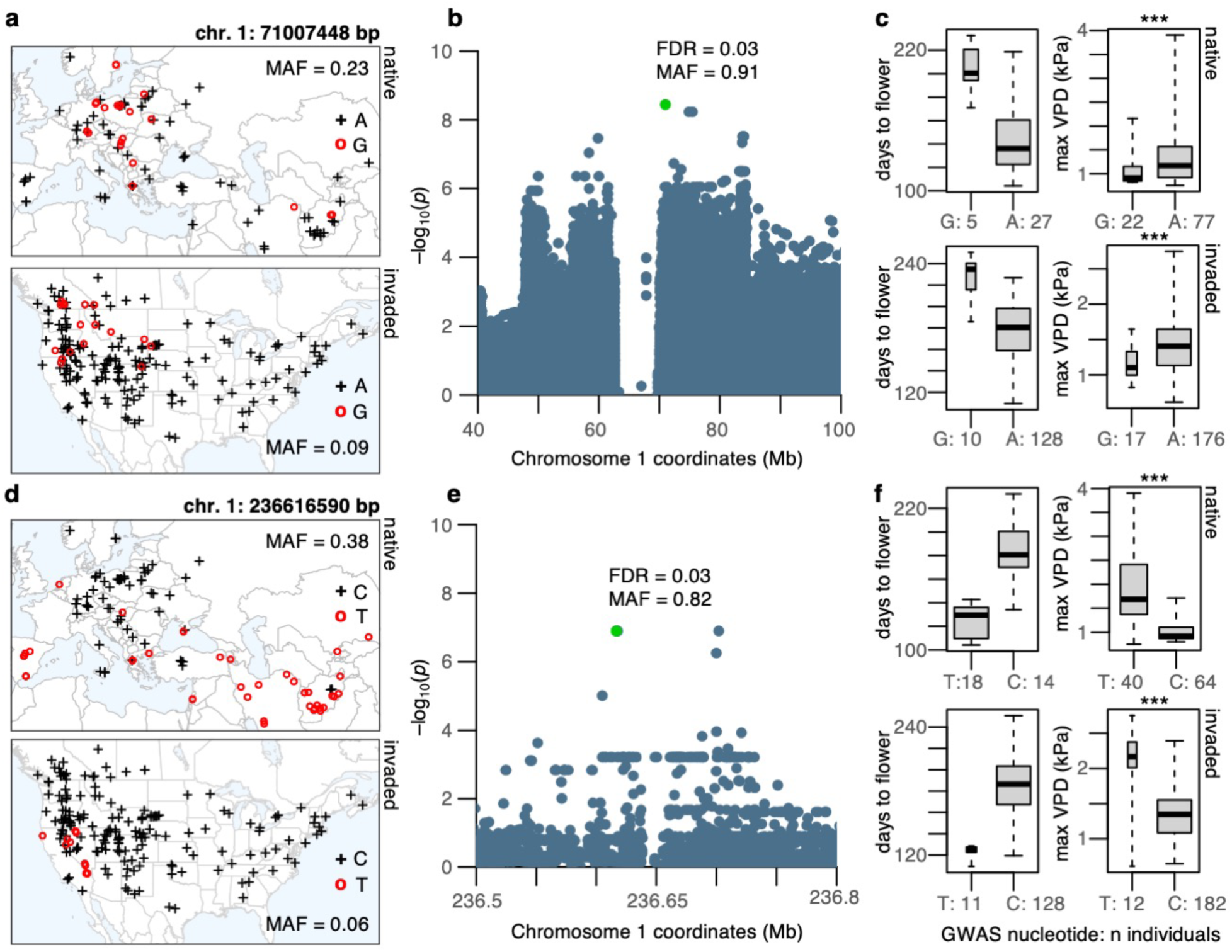
Environmental trends of two flowering time QTL are mirrored between native and invasive genotypes. (**a** and **d**) Geographic distribution of QTL SNP alleles in the native (top) and invaded (bottom) range; crosses represent the reference/major allele, and open circles the alternate/minor allele. (**b** and **e**) Zoomed-in Manhattan plots showing Wald-test *p*-values (plotted as –log_10_) from GWAS and genomic location of top SNP (marked in green), with respective false-discovery-rate (FDR) and minor allele frequency (MAF). (**c** and **f**) Phenotypic (boxplots to the left) and environmental variation (boxplots to the right) of flowering time QTL SNP alleles identified with GWAS. Boxplots indicate median (middle line), 25^th^, 75^th^ percentile (box), and whiskers cover the data extent. \*\*\**p*<0.0008 from two-tailed t-tests, but kinship linear-mixed models showed no significant differences. Max VPD: Maximum vapor pressure deficit in kPa. A, G, T, C on maps and x-axis of boxplots indicate nucleotides.

We annotated SNPs based on genome-wide linkage-disequilibrium (LD, Supplementary Fig. 12a). We found that at 194.5 kb, LD decayed to ∼80% of the background LD (here taken as 5 Mb). A similar LD decay pattern was observed across chromosomes (Supplementary Fig. 12b), while between chromosomes average R^2^ ∼0.1. Below, we thus highlight QTL based on the position of the closest gene within a 200 kb window centered at the GWAS SNP. We focus on flowering time because this was the only trait for which we found gene functions clearly related to phenotype.

The top flowering time QTL (detected with LMM) contained multiple SNPs along a haploblock of ∼28 Mb (chromosome 1: 56–84 Mb, allele frequency (AF)∼0.9) containing ∼64 genes with annotations based on homology to *Oryza sativa* and *Arabidopsis thaliana*. These genes were enriched for gene ontology terms describing developmental processes involving reproductive structure/system, embryo, embryo ending in seed dormancy, post-embryonic, fruit, and seed (8 *O. sativa* genes and 14 *A. thaliana* genes, *p*<0.0003, FDR=0.01). Such a large haploblock could indicate a structural variant, a potential driver of local adaptation^51,52^. This locus thus merits further investigation.

The top SNP of the haploblock (chromosome 1: 71007448 bp, AF=0.91; top in LMM and 2^nd^ top in MLMM) was 25 kb downstream of a *O. sativa* homolog, the DnaJ protein Erdj3b. Expression of Erdj3b in *O. sativa* is critical for heat stress tolerance during seed development^53^. Late flowering alleles were more frequent in humid/colder regions of the native (two-tailed t-test t= – 3.66, df=83.61, *p*=0.0004) and invaded range (two-tailed t-test t= –4.31, df=26.41, *p*=0.0002) (Fig. 4a–c), suggesting cheatgrass adaptation to temperature gradients may be linked to seed sensitivity to temperature stress.

The fourth top flowering time QTL (only found with LMM) comprised three SNPs (chromosome 1: 236616590 bp, 236616999 bp, 236617691 bp, AF=0.82) 0.5 kb upstream (putative promoter region) of the *A. thaliana* homolog *ATE1* (AT5G05700). *ATE1* regulates seed maturation, seedling metabolism, and abscisic acid germination sensitivity^54^. Early flowering alleles were found in drier regions of the native (two-tailed t-test t=6.73, df=42.19, *p*=3.4e^−8^) and invaded range (two-tailed t-test t= 4.82, df=11.67, *p*=0.0004), specifically the Mediterranean and west Asia in the native range, and the Mojave and Lahontan Basin in the invaded range, but also reaching Mediterranean climates of coastal WNA (Fig. 4d–f). These patterns suggest that even the specific QTL underlying local adaptation in the native range have been similarly reused for local adaptation in the invaded range.

We compared our GWAS results with a published study that performed a GWAS for flowering time using a smaller and much less diverse genotype panel^32^. There was no overlap among the 200 top GWAS SNPs we found (100 top SNPs × 2 GWAS methods) and the SNPs detected in that study (Table 2 in^32^), likely due to our larger and more diverse panel of genotypes.

### Cheatgrass dominates where local adaptation is predicted to be stronger

Whole genome-environment associations in the native range predicted local adaptation in the invaded range, especially where cheatgrass is most dominant. To further evaluate whether invasive genotypes matched local climates as in the native range, we used a predictive genome-environment model. Using the native range RDA model of genotype as a function of climate (Fig. 2e), we first predicted invasive genotypes for locations of our sequenced samples. Next, we calculated the genetic distance between predicted and observed genotypes, similar to metrics sometimes referred to as ‘genomic offset’^55^. Genotype-environment matching (*i.e.*, low genetic distance, or offset, between predicted and observed genotypes) was strongest at northern latitudes across North America, particularly in WNA. Putative maladaptation (*i.e.*, high genetic distance between predicted and observed genotypes) was strongest in the southeast USA (Fig. 5a). By comparing mean genetic distance to means of 1000 null permutations, we found the mean genetic distance was significantly lower than the null expectation in WNA (*p*<0.002), but not in ENA (*p*=0.5, Fig. 5b). This finding is consistent with the hypothesis that local adaptation to climate in WNA reflects the patterns observed in the native range, while cheatgrass in ENA has a novel strategy or is not locally adapted. Unlike WNA, cheatgrass populations in ENA are more restricted to highly disturbed urban and agricultural sites, rarely forming large monospecific stands^56^.

**Fig. 5:**
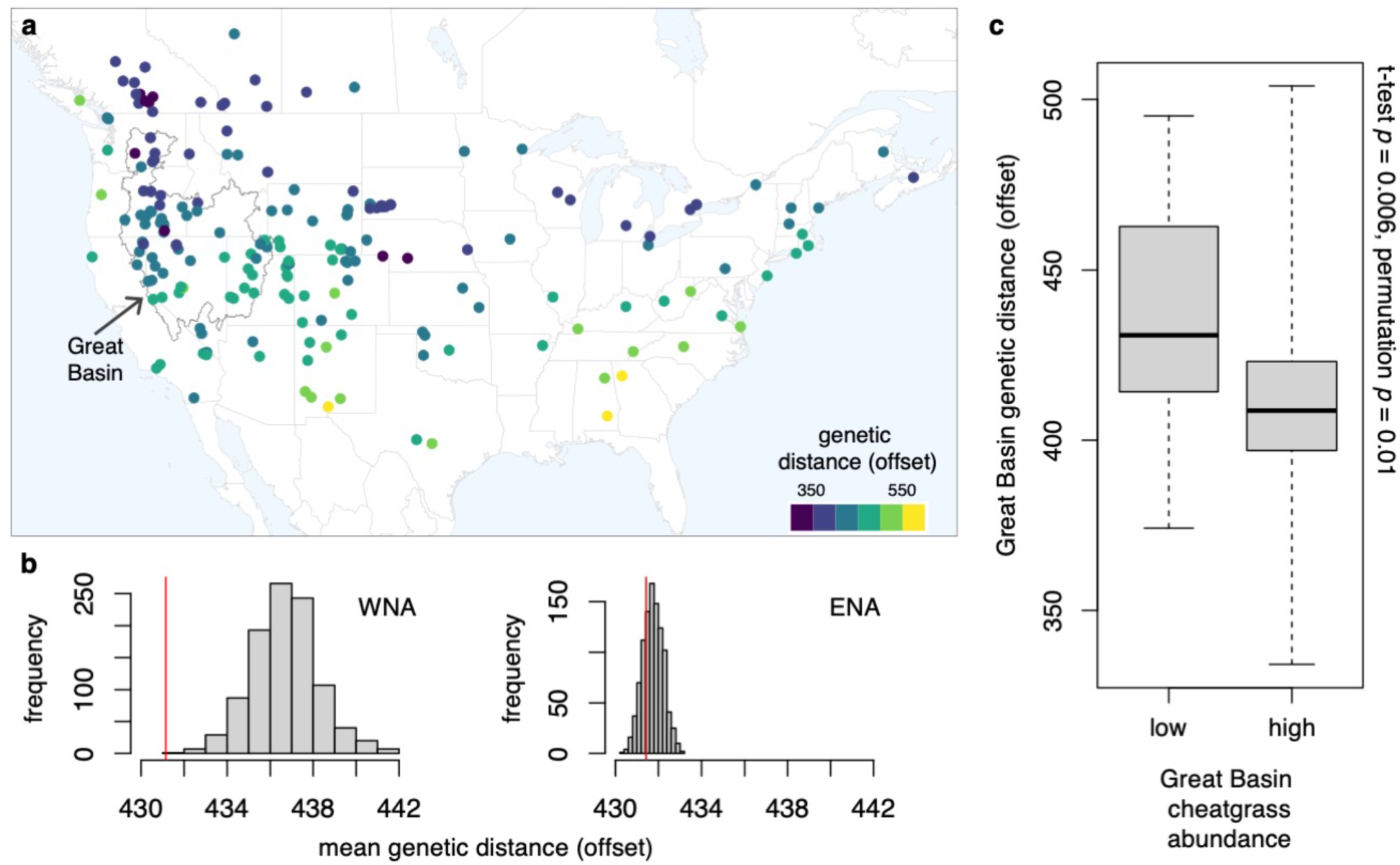
Genomic predictions of strong local adaptation occur in regions where cheatgrass is most dominant. (**a**) Geographic distribution of the genomic offset estimated for each invasive genotype (n=194). The genomic offset or maladaptation is the genetic distance between observed invasive genotypes and the genotype-environment predictions in the invaded range based on the native range genotype-environment association. (**b**) Histograms of the mean genetic distance (offset) of 1000 null permutations in western North America (WNA, n=127) and eastern North America (ENA, n=67), relative to their estimated mean genetic distance (red lines). (**c**) Within the Great Basin (polygon in a, n=55), the mean genetic distance (offset) is significantly lower in areas where cheatgrass occurs in high (*i.e.*, representing >15% vegetation cover) vs. low abundance. Boxplots indicate median (middle line), 25^th^, 75^th^ percentile (box), and whiskers cover the data extent.

To assess whether matching of specific genotypes to local environments promotes cheatgrass invasion, we compared the strength of genotype-environment correlations with variation in cheatgrass abundance from 11,307 field surveys across the Great Basin (Fig. 5a), the region where the invasion has its worst impacts^30^. Locations where cheatgrass occurs in high abundance showed significantly high genotype-environment matching based on the native range model compared to sites where cheatgrass does not dominate (n=55, two-tailed t-test t=2.89, df=52.3, *p*=0.006, Fig. 5c), suggesting local adaptation promotes cheatgrass dominance. This pattern was consistent when comparing genotype-environment matching of high-abundance sites to 1000 null permutations of genotypes within the Great Basin (*p*=0.01), evidence that this pattern was not merely due to environmental characteristics of the low-abundance sites but reflects the match of genotypes to their local environments.

### Synthesis

Biological invasions pose a major environmental threat, but the roles of genomic diversity, repeated introductions, and adaptation are poorly understood. Our results have important implications for understanding the evolution of local adaptation in invasive species that have greatly expanded their range. Past studies suggest that selfing species with large native ranges – like cheatgrass– are more likely to establish self-sustaining populations in new regions^23,57,58^, but the mechanisms promoting successful establishment have been less explored.

We show that multiple diverse introductions and long-distance dispersal post-introduction likely increased the chances of cheatgrass genotypes arriving to favorable North American environments. Across Eurasia, cheatgrass shows clines indicative of local adaptation and continuous isolation by distance, likely shaped by multiple migrations and subsequent isolation after the Last Glacial Maximum^45^. In the invaded range, many genotypes represent a mosaic of near-clonal lineages across sometimes widely spread locations but similar environments, and non-clonal genotypes sorted along the steep climate gradients of western North America. Despite differences in genomic diversity and likely demographic history between the native and invaded ranges, western North American genotypes closely matched the native signatures of local adaptation along temperature/aridity gradients. Thus, in North America environmental filtering of pre-adapted genotypes likely led to reuse of native diversity and facilitated range expansion^18,19,59^. Accordingly, common gardens revealed that changing selection likely maintains a major life history cline in the USA Intermountain West. Furthermore, genomic signatures of local adaptation also predicted cheatgrass ecological dominance across the USA Great Basin, indicating that factors supporting local adaptation, such as high genetic diversity from repeated introductions, fuel the invasion.

Our findings emphasize that any sources of genetic diversity could continue to reshape adaptation in established invasive species^60^. With high genomic diversity and no dispersal limitation, range-wide adaptation could persist over time even under shifting environments. Limiting ongoing introductions and intra-continental dispersal of genotypes (*e.g.*, by limiting seed contaminants in grains) could likely help minimize rapid adaptation of invasive plants. For annual selfers like cheatgrass, this strategy might limit local adaptation via pre-adaptation, but also via *de novo* variation from uncommon but potentially important outcrossing events^28,39,47^.

## METHODS

### Plant material

Natural inbred lines of *Bromus tectorum* were obtained from 1) the Genome Resources Information Network (GRIN), 2) Greenhouse inbred/selfed plants (S1 or S2) of field samples in western North America collected 2019–2020, 3) field samples from the native and invaded range collected 2020–2022, and 4) DNA extractions of frozen seedlings (contributed by Brian Rector, USDA–ARS) for 29, 111, 155, and 12 samples, respectively (Supplementary Data 1). With this panel of genotypes, we targeted sites with distinct environmental conditions, favoring environmental variation over intra-population sampling. Sites were ∼1–6600 km apart within the native or invaded ranges: 194 North American, 105 Eurasian, and 8 from regions with less extensive invasions: 2 from Argentina, 1 from Australia, 3 from New Zealand, and 2 from South Korea. Genotypes with available seeds (295) were germinated in a growth chamber at 20°C (80% humidity, 12 h light/12 h dark, 200 μmol m^2^ s^-1^ light intensity) to increase seed (Supplementary Methods 1), verify identification, and obtain tissue for whole genome sequencing (Supplementary Methods 2–4) and genotyping (Supplementary Methods 5–6).

Native genotypes were assigned to central-north-east Europe, Mediterranean, or west Asia based on geographic location (Supplementary Data 1). North American genotypes were assigned to eastern North America (ENA) or western North America (WNA) based on ecological region at location of origin (Supplementary Methods 7). WNA: marine west coast forest, Mediterranean California, North American deserts, northwestern forested mountains, and temperate Sierras. ENA: eastern temperate forests, Great Plains, and northern forests (Supplementary Fig. 2).

### Population genetic structure

With a dataset of 266,504 unlinked sites, we used multiple methods to infer population genetic structure that we interpret collectively (Supplementary Methods 8–9). We estimated individual admixture proportions in NGSadmix (v.33)^61^ and inferred population genetic structure with PCA in PCAngsd (v.1.11)^62^, which work directly with genotype likelihoods that contain all relevant information of unobserved genotypes. Individual admixture proportions were estimated with maximum likelihood in 12 replicates, for K=2–12 genetic clusters, on sites with minor allele frequency (MAF) >0.05, at least in 50% individuals, and <75% missing data. Cross-validation of number of clusters was determined from log-likelihoods of the NGSadmix output across all replicates^63^.

We also computed an unrooted phylogenetic tree with the Neighbor-Joining (NJ) algorithm^64^ and calculated Nei’s^65^ pairwise F_ST_ and Weir & Goudet’s population-specific F ^66^ based on genetic dissimilarity estimated with SNPRelate (v.0.9.19)^67^. Pairwise F_ST_ measures population genetic differentiation between regions, whereas population-specific F_ST_ measures regional deviations from the ancestral population. High values of population specific-F_ST_ indicate high within-group allele sharing and potentially greater divergence from ancestral populations, while low values indicate possible ancestral populations.

### Genomic diversity and Tajima’s D

We compared nucleotide diversity (π^68^) between ranges and regions in the native and invaded range using a dataset of 15,101,725 sites (Supplementary Methods 6). Regional .vcf datasets containing all sites were generated with BCFtools (v.1.18)^69^ (view -S), and genome-wide estimates of π were obtained with VCFtools (v.0.1.15)^70^ using 50 kb sampling windows. Deviations from neutral evolution between geographic regions were examined with the Tajima’s D statistic^71^, which compares the mean number of pairwise differences against the number of segregating sites observed in a set of sequences. Tajima’s D for each region was calculated in 50 kb sampling windows for shared SNPs in VCFtools using -- TajimaD.

### Genetic load

Genotype mutation load (under the hypothesis that most protein changing mutations are deleterious) was estimated separately from the high impact and missense variants, both normalized by (divided by) the number of synonymous variants (Supplementary Methods 10). We used 2-way ANOVA and Tukey HSD tests to examine differences in genetic loads between native and invaded range genotypes of the same ancestry. Genotypes were assigned to a cluster based on having >0.55 ancestry proportion for the NGSadmix K=4 ancestral genetic clusters. If no K=4 ancestry was >0.55, genotypes were designated as intermediate.

### Self-fertilization rates

We examined the causes of inbreeding between native and invaded genotypes with several statistics. We used a subset of 101 closely related native and invasive genotypes that were sequenced from seedlings of plants collected directly from the field (as opposed to a greenhouse bulking or GRIN). Based on the ∼15M SNPs dataset, we first computed runs of homozygosity^46,72^ (with BCFtools roh), Tajima’s D (with VCFtools --TajimaD in 100 kb windows), and heterozygosity (with PLINK (v.1.9)^73^ --het) for one lineage represented in the native and North American invaded range (native n=27, invaded n=74). Then we implemented random forests in R to analyze these genetic statistics together and estimate selfing rates for each group using a recently published model (“sequential model”^46^). We compared group means with a two-tailed t-test.

### Isolation-by-distance

We examined isolation-by-distance in native and invasive genotypes, and in genotypes from WNA and ENA, with the same LD-filtered SNP dataset. Genome-wide pairwise genetic dissimilarity matrices were obtained as described above. Pairwise geographic distances were calculated in kilometers from genotype coordinates with the spDists function in the R package sp (v.2.1-3)^74^, using the WGS84 ellipsoid projection. Simple Mantel tests^75^ were used to test if the natural logarithm of geographic distance predicts genetic distance with the function mantel.rtest in the R package ade4 (v.1.7-22)^76^, using 999 permutations. We conducted linear regressions (with the R function lm) to assess the proportion of genetic variance explained by geographic distance. We also used the mantel.rtest R function to assess how climatic distance changed with geographic distance.

### Environmental differentiation between groups and across space

We examined if the 19 genetically different groups of 2–14 nearly clonal genotypes (detected based on >99% SNP similarity) were differentiated by environment using climate data at their location of origin. We used clonal group identity as the predictor of 52 CHELSA climate variables^77,78^ (Supplementary Methods 7, Supplementary Data 1, Supplementary Fig. 7b) with PERMANOVA using function adonis2 (9999 permutations, Euclidian distances) in the R package vegan (v.2.6-4)^79^. To examine spatial environmental heterogeneity, climatic distance was estimated for each region (native, ENA, WNA) based on pairwise Euclidean distances in the environmental PCA produced above, using the function vegdist in vegan.

### Variance partitioning of genomic diversity

Redundancy analyses (RDA) were used to model how sets of variables explained SNP variation and for identifying abiotic gradients explaining the most genome-wide SNP variation^80^. To model geographic patterns in the RDA, a distance matrix obtained from coordinates was converted into a spatial weighting matrix to get a reduced-dimension set of orthogonal variables (Moran’s eigenvector maps, MEMs^81^). MEMs are eigenvectors of the pairwise spatial weighting matrix among samples (Supplementary Methods 11). Then, RDA was conducted with variance partitioning^80^ to quantify proportion of genome-wide SNP variation explained by each of two categories of covariates: Abiotic variables and geographic MEMs. We selected abiotic variables that were informative and non-colinear based on the PCA explaining range-wide environmental variation (Supplementary Fig. 6e). Variance partitioning estimates proportion of SNP variation that is explained by the collection of variables in each category and by collinearity among variables. To identify environmental gradients associated with genome-wide divergence, RDA was also conducted using only abiotic variables for native and invasive genotypes. We computed RDA and performed variance partitioning with functions rda and varpart, respectively, in vegan.

### Ancestry-environment associations

To assess environmental filtering of pre-adapted genotypes in North America, we examined ancestry-climate associations in invasive versus native genotypes using generalized additive models (GAMs). GAMs allow us to account for nonlinear patterns between predictors and the response variable^82^. Environmental predictors were the same aridity and temperature gradients used for trait-environment clines, in addition to precipitation seasonality (all representative of climatic variation in cheatgrass genotypes; Supplementary Fig. 6e). For each NGSadmix ancestral cluster (K=4), GAMs were implemented with the function gam in the R package mgcv (v.1.9-1)^83^ with a logit link function and beta-distributed residuals. Genotypes were assigned to a cluster based on having >0.55 ancestry proportion for the NGSadmix K=4 ancestral genetic clusters. Intermediate genotypes *(i.e.*, composed of multiple ancestries) were excluded from this analysis.

### Phenotypes

During the grow out in 2020, we measured phenotypes on up to 184 genotypes with 2–3 replicates that emerged within ∼9–18 days of planting and survived until harvesting. Eleven phenotypes were recorded: seedling and adult (*i.e.*, reproductive) height (used to get spring growth), number of leaves, number of tillers, days to flower, inflorescence height, dry biomass, total seed mass (*i.e.*, fecundity), individual seed mass (*i.e.*, seed mass), total seed length, and awn length (Supplementary Methods 12).

After quality/error checking, the best linear unbiased estimate (BLUE) of phenotypes was calculated per genotype with the BLUE function in the R package polyqtlR (v.0.1.1)^84^, using genotype as the predictor of trait measurements across 2–3 replicates and tray as a random effect. We then calculated broad sense heritability (H^2^) of traits as the proportion of phenotypic variance explained by genotype in a linear model. The total set of phenotyped genotypes included: 184 with vegetative height/growth/count data, 173 with flowering/inflorescence data, 178 with dry biomass data, and 182 with seed data, for a total of 169 genotypes with no missing phenotypes.

### Trait variation and environmental associations

To detect axes of life history variation, we summarized the natural genetic variation in our growth chamber phenotypes with PCA (function prcomp, variables scaled and centered) in R. PC1, PC2, and flowering time, were then used as response variables for investigating trait differences between ranges and environmental gradients in phenotypes. To assess differences in trait means between native and invasive genotypes we implemented two-tailed t-tests as well as linear mixed models that accounted for kinship between genotypes (see below). To assess phenotypic differentiation between groups of nearly clonal genotypes, we performed PERMANOVA with group identity as the predictor and eleven phenotypes as response with function adonis2 in the R package vegan (9999 permutations, Euclidian distances). To assess trait-environment clines we used kinship linear-mixed models, which when significant (*i.e.*, p≤0.05), they provide evidence of selection (vs. population genetic structure/drift) explaining variation, similar to Q_ST_–F_ST_ tests^49^. A kinship matrix was estimated with identity-by-state (IBS, *i.e.*, allele sharing between pairs of genotypes) using the dataset of 15,101,725 SNPs and function snpgdsIBS in the R package SNPRelate. This kinship matrix was used to fit linear mixed models with random genotype effects using function lmekin in the R package coxme (v.2.2-20)^85^. We focused on maximum monthly vapor pressure deficit (Pa), describing aridity, and mean air temperature of the coldest quarter (°C), describing winter temperature. These two climate variables showed the highest loads on PC1 and PC2, respectively, on a PCA of 52 climate variables for the 307 native and invasive genotypes (Supplementary Fig. 6e). To test if clines were repeated, absent, or shifted in the invaded relative to the native range, we also tested for an interaction between ranges in the models.

### Field common gardens

We conducted a replicated common garden experiment with cheatgrass genotypes in the 2022 and 2023 growing seasons, across two sites in the Intermountain West that varied in their regional climatic conditions: a cool site with little temperature seasonality (Sheep Station, ID [44.2456°N, 112.2144°W]) and a warm site with pronounced temperature seasonality (Wildcat, ID [43.4744°N, 116.9018°W]). We grew replicates of 95 genotypes from Fall 2021 to Spring 2022 and 93 genotypes from Fall 2022 to Spring 2023 at two different densities (low=100 seeds/1 m^2^; high=100 seeds/0.04 m^2^) and under two different temperature treatments (low=white gravel; high=black gravel) in a factorial design at both sites^50^ (Supplementary Methods 13).

We compared the direction and magnitude of selection on flowering phenology between the cold, less seasonal site and the warm, seasonal site for both the 2022 and 2023 growing seasons. For each year and common garden site combination, we calculated the average fitness (2022=seed count; 2023=reproductive biomass) and average first flowering day for each genotype, across all treatments. In the calculations of average first flowering day, if a plant did not flower (*i.e.*, fitness=0), it was assigned the average first flowering day for all plants of that genotype that did flower at some point during the growing season. For each growing season year, we regressed the mean fitness data on the mean flowering time data across genotypes and compared the slopes between the cold, less seasonal site and the warm, seasonal site. Positive slopes on this graph indicate that flowering later is selected, while negative slopes indicate that flowering earlier is selected. Models that included a random intercept for each genotype, with a correlation structure specified by a kinship matrix, also provided evidence of selection. To confirm that our larger dataset on flowering time from a growth chamber was consistent with field measurements, we compared these values and found them correlated with Pearson r=0.6.

### Genome wide association studies (GWAS)

To identify QTL for growth chamber phenotypes, we implemented GWAS that controlled for kinship on our BLUEs dataset (n=173–184 genotypes, 14.6–14.7M SNPs excluding SNPs with MAF<0.05) using two methods: a univariate linear mixed model (LMM) and a multilocus mixed model (MLMM). SNP genotype data were generated with function snpgdsGetGeno in the R package SNPRelate. The univariate LMM was fit with gemma (v.0.98.5)^86^ with an IBS matrix as the random effect accounting for relatedness between genotypes. The MLMM was implemented with FarmCPUpp^87^, which computes a restricted kinship matrix based on pseudo-quantitative trait nucleotides (QTN) selected from a preliminary GWAS step. FarmCPUpp also takes principal components of genome-wide SNP variation as covariates (here PC1–PC3), providing a stronger control of population genetic structure. Principal components were obtained with function snpgdsPCA in the R package SNPRelate. Moreover, while gemma tests for an association with each SNP individually, FarmCPUpp performs additional steps that detect pseudo-QTNs based on LD and uses model selection and multiple regression to retain the best set of pseudo-QTNs. Thus, while gemma might reveal large blocks of significantly associated SNPs (which might correlate with chromosomal rearrangements), this signal should be lost with FarmCPUpp (which might be better at detecting causal SNPs). To detect statistical significance of GWAS SNPs, we used a false-discovery-rate (FDR) threshold of 0.05 on output *p*-values. Associations were inspected with Manhattan plots and model fits were assessed with quantile-quantile (Q-Q) plots.

### Linkage disequilibrium (LD) and SNP annotation

To assign GWAS SNPs to annotated cheatgrass genes, we first investigated linkage disequilibrium (LD) decay with genomic distance in our sequenced panel of 307 genotypes. We used PopLDdecay^88^, which takes a .vcf with samples and computes the square of Pearson correlations (R^2^) between pairs of SNPs genome-wide (using the ∼15M SNPs dataset) or per chromosome. We excluded SNPs with MAF<0.05 and >5% heterozygote individuals and calculated R^2^ between pairs of SNPs at a minimum of 10 bp apart and a maximum of 5 Mb apart. We detected the pattern of LD decay by plotting mean LD values in 100 bp bins from 0–5Mb genomic distance. We observed substantial long-range LD with mean R^2^∼0.3 even at 5Mb (Supplementary Fig. 12a), in line with strong population structure. However, there was a clear decay in LD with genomic distance. The initial mean LD (R^2^=0.45) at 10 bp decayed halfway (R^2^=0.376) of the minimum observed LD (R^2^=0.3) by 194.5 kb. Because mean LD was high even at 5Mb, we also assessed inter-chromosomal LD (Supplementary Methods 14).

### QTL-environment clines and enrichment analysis

Environmental variation of allele frequency in QTL detected with GWAS was examined with two-tailed t-tests and kinship linear-mixed models. QTL-environment tests were performed separately for the native and invaded range. To find significantly over-represented GO terms or parents of these terms in the haploblock detected for flowering time, we uploaded a gene set to the PlantRegMap GO Term Enrichment tool^89^, based on *A. thaliana* and *O. sativa* homologs obtained from CoGe annotations.

### Genome-environment matching of invasive genotypes

We further tested for evidence of pre-adaptation by quantifying how well a native range genotype-environment association (GEA) model predicted genetic composition in the invaded range. Our approach is like the genomic offset statistics used for predicting climate change impacts on maladaptation^55,80^. We used the RDA-generated native range GEA to predict maladaptation of invaded range genotypes, separately for WNA and ENA, with the function ‘predict’ in the R package vegan. The Euclidian distance between predicted and observed allele frequencies for each invasive genotype was then estimated, representing the genetic maladaptation to the invaded range site. To evaluate if the mean genetic maladaptation WNA and ENA was different from random, predictions and genetic (*i.e.*, Euclidean) distances were recalculated in 1000 reshuffled environments. Genetic maladaptation was considered significantly lower than expected by chance if it fell in the lower 0.025 tail of the random distribution of mean genomic distances in a two-tailed test.

### Genome-environment matching and invasive spread

We examined if cheatgrass dominance was correlated with the strength of local adaptation using available cheatgrass abundance data. Using random forest models, Bradley *et al.*^30^ created a regional classification of cheatgrass presence across the Great Basin based on 11,307 surveyed sites. Their classification differentiates between cheatgrass present at high abundance (≥ 15%) and cheatgrass absent or at low abundance (< 15%). Genetic maladaptation/offset was compared between sites where cheatgrass is in high abundance to sites where cheatgrass is in low abundance (high vs. low in Fig. 5c, respectively) with a two-tailed t-test (n=54 sites or Great Basin genotypes). To assess if this pattern was driven by the match of local genotypes to local environments (as opposed to the environmental characteristics of the low abundance cheatgrass sites), we recalculated genomic offset in 1000 reshuffled Great Basin environments and compared offset of regions where cheatgrass dominates to the 1000 null permutations of genotypes within the Great Basin.

## Data availability

Seeds of *Bromus tectorum* genotypes used in this study are available from the lead contact upon request and will be deposited to GRIN. Herbarium vouchers of genotypes will be deposited at The Pennsylvania State University Herbarium (PAC). Whole-genome sequences for 303 genotypes are being deposited in the European Nucleotide Archive (ENA; https://www.ebi.ac.uk/ena). Whole-genome sequences for four genotypes sequenced by the DOE Joint Genome Institute are publicly available at their website under proposal ID: 506608. The beagle (genotype-likelihoods), vcf (SNP calls), and raw phenotype data will be publicly available at Figshare^90^. Genotype-level geographic information, climate data, growth chamber phenotypes, ancestry proportions, GWAS SNPs, and field common gardens data are in Supplementary Data 1. Gene annotations of GWAS results (100 top SNPs) are in Supplementary Data 2.

## Code availability

Code used to analyze NGS data will be publicly available at Figshare^90^.

## Acknowledgments

Any opinions, findings, conclusions, or recommendations expressed in the material are those of the authors and should not be construed to represent any official USDA determination or policy. Any use of trade, firm, or product names is for descriptive purposes only and does not imply endorsement by the U.S. Government. Xavier Mack, Carlos Rodríguez-Gonzalez, Yuxin Luo, and Katherine Blocklove helped with measuring phenotypes, harvesting, and performing DNA extractions. Ian Burke, Samuel Revolinski, Peter Maughan, and Craig Coleman granted us access to the reference genome prior to publication. The USGS FIREss team contributed to common garden establishment and data collection on the Wildcat site. iNaturalist was a key resource for identifying seed collectors, among those were also Dave Barnett, David Board, Chalon Boesel, John Bradford, Jaime Braschi, Howard Bruner, Victoria Bustamante, Charles Campbell, Jeanne Chambers, Mike Chen, Mark Chynoweth, Jason Cooper, Massimo Cristofaro, Melodie Cunningham, Kirk Davies, Janelle Downs, Torsten Eriksson, Maggie Eshleman, Erica Fleishman, Carol Kadonsky & Bill Foreman, Berit Gehrke, Tom Getts, Richard Gill, Dana Hartel, Nate Hartley, Patricia Hollins, Alex Hood, Tayla Hook, Parker Hopkins, Becky Hufft, Jennifer Kalt, Lorri Kendrick, Molly Ladd, Matt Lavin, Steven Lee, Jonathan Levine, Marisa Mancillas, John Maron, Grace McCartha, Randal Mindell, Chandra Moffat, Brooke Moore, James Nagler, Yael Orgad, Matthew Pedrotti, David Pyke, Sasha Reed, Matt Rinella, Viktoria Ropak, Håkan Rydin, Geno Schupp, Adam Searcy, Tim Springer, Amy Symstad, Tracy Thomas, Trudy Trevarthen, Samantha vanDeurs, Biljana Vidovic, Viktoria Wagner, Gretchen Whetham, David Wilderman, Eileen Wyza, and Pauline & Jon Zweck. Funding: National Science Foundation grant DEB-1927282 (PBA), DEB-1927009 (JRL), DEB-1927177 (MBH); Joint Genome Institute of the U.S. Department of Energy grant New Investigator Award-506608 (JRL); National Institutes of Health grant R35GM138300 (JRL).

## Author contributions

Conceptualization: DG, CSB, EAL, DMB, MJG, LMP, MBH, PBA, JRL

Methodology: DG, TMM, NP, SR, AP, CSB, EAL, DMB, MJG, LMP, MBH, PBA, JRL

Investigation: DG, MLV, AD, RB-Z, EAL, DMB, MJG, LMP, PBA, JRL, OB, CSB, RB, SB-M, SMC, SLD-F, AD, RCD, PWD, DJE, TAV, HH, MCH, RAH, SJ, JMK, EK, SK, MK, AK, ML, SL, DM, CLM, JWM, DUN, JPO, RP, LAP, LR, BGR, CR, MS, RKS, AS, BMS, RLS, KGT, AKU, AVW, C-AW, JZ.

Funding acquisition: PBA, MBH, JRL, MJG

Project administration: PBA, MBH, CSB, DMB, MJG, LMP, JRL

Writing – original draft: DG, JRL Writing – review & editing: all authors

## Additional information

**Supplementary information** is available for this paper.

**Correspondence** and requests for materials should be addressed to D.G.

**Reprints and permission information** is available at http://www.nature.com/reprints

