## Supplementary for "Local adaptation to climate facilitates a global invasion"

**This PDF file includes**:

Supplementary Methods 1 to 14

Supplementary Table 1

Supplementary Figures 1 to 12

Supplementary References

**Supplementary Methods**

Citations throughout these sections are included in the References of the main text.

**1. Growth conditions**: The first seed bulking was performed in 2020–2021 (193 genotypes) and the second in 2022 (102 genotypes). Three replicates were grown in a 1:1 mix of commercial grade sand and growing medium (PGX PRO-MIX, Premier Tech Manufacturer) to obtain seeds. Two replicates were grown in 100% growing medium to obtain tissue for DNA. Plants were grown in conetainers (1.5-inch diameter, 164 ml) in a RL98 rack (Stuewe & Sons, Inc.). Cotton balls were placed at the bottom of each conetainer to prevent soil loss through drainage holes. Plants selected for DNA extraction were kept under constant temperature until tissue collection. Seedlings (~5–15 day-old plants) of the grow out sets were vernalized in a cold room at 4ºC (30% humidity, 8 h light/16 h dark, ~75 µmol $m^2$ $s^{-1}$ light intensity) for 10 weeks, then transferred back to the growth chamber at 20ºC day/15ºC night (50% humidity, 14 h light/10 h dark, 200 µmol $m^2$ $s^{-1}$ light intensity) and kept well-watered until harvesting. Plants were randomized within trays, with every other position left empty within racks (for 49 pots/tray), and once plants flowered, surrounded with a hard plastic transparent cylinder (with air-flow holes) attached to the base of pots to avoid any outcrossing. Trays were periodically rotated (twice per week) and moved around the growth chamber (twice per month) to mitigate positional effects.

**2. DNA purification**: Total genomic DNA for each genotype was extracted from ~100 mg of fresh tissue collected from one healthy individual in the DNA plant set, using the Viogene Plant Genomic DNA Extraction System (Viogene BioTek, Corp.). To ensure purity and high molecular weight, DNA was subsequently cleaned with 0.9X AMPure XP magnetic beads (Beckman Coulter, Inc.). DNA purity was assessed with a Thermo Scientific NanoDrop 2000 spectrophotometer (Thermo Fisher Scientific, Inc.) and quantified with an Invitrogen Qubit 2.0 fluorometer (Thermo Fisher Scientific, Inc.).

**3. Next generation whole-genome sequencing**: A set of 303 genotypes were sequenced at an average coverage of ~1.2–4.8X per genotype (genome size ~2.5 Gb) at the Texas A&M AgriLife Research. At least 500 ng of genomic DNA was used to construct paired-end sequencing libraries (PerkinElmer NEXTFLEX Rapid XP DNA-Seq Kit HT) which were sequenced on an Illumina

NovaSeq 6000 S4 platform - 2x150 v.1.5 (Illumina, Inc.). Sequence cluster identification, quality prefiltering, base calling and uncertainty assessment were done in real time using Illumina's NCS 1.0.2 and RFV 1.0.2 software with default parameter settings. Sequencer .cbcl basecall files were demultiplexed and formatted into .fastq files using bcl2fastq 2 2.19.0 script configureBclToFastq.pl. Raw reads were processed with FastQC (v.0.11.8)[1] and filtered for low quality and adapter regions using Trimmomatic (v.0.39)[2]. Filtered .fastq files contained ~20–80 × $10^6$ reads totaling ~3–12 Gb per genotype.

A separate set of four genotypes was sequenced at an average of 70–120X per genotype at the Joint Genome Institute for genome size estimation. At least 10 µg of genomic DNA was used to construct libraries for Illumina sequencing as above. Duplicate reads were removed based on paired sequence matching using Clumpify[3]. BBDuk (v.38.90)[3] was used to trim reads that contained adapter sequences and homopolymers of G's of size ≥5 at the ends of reads. BBDuk was also used to remove reads that contained one or more 'N' bases, had an average quality score <6, or had a minimum length ≤49 bp or 33% of the full read length. Filtered .fastq files contained ~1.16–2.04 M reads totaling ~174–308 Gb per genotype.

**4. Mapping reads to the reference genome**: Paired-end reads from each genotype were mapped to the *Bromus tectorum* genome using BWA-MEM (v.0.7.15)[4] with default parameter settings. Then in SAMtools (v.1.18)[5] read alignments were converted into BAM format (with: view -bhS), sorted by read names (sort -n) to check and update mate coordinates (fixmate -rpcm), sorted by genomic coordinates (sort) to mark and remove PCR duplicates (markdup -Srs), and filtered for incomplete and poor quality alignments (view -b -m 30 -q 30).

**5. Initial variant calling**: Analysis of Next Generation Sequencing Data (ANGSD v. 0.938)[6] software was used to detect SNPs and calculate genotype likelihoods across the 307 samples, thus allowing to account for genotype uncertainty in downstream analyses. An initial variant calling step was performed on all samples, separately for each chromosome, using base call and mapping quality filters (-uniqueOnly 1 -minMapQ 30 -C 50 -baq 1 -minQ 20 -remove_bads 1 -only_proper_pairs 1) with the SAMtools genotype likelihood framework (GL -1) and the output written to Beagle format (-doGlf 2). To avoid potential biases arising from sequencing errors and excessive repetitive regions, we kept biallelic SNPs with a sequencing coverage of minimum 5× and maximum 25× in 50% of all samples and with an allele frequency above 0.05 (-SNP_pval 1e-6 -doCounts 1 -setMinDepth 1535 -setMaxDepth 7675 -minInd 154 -minMaf 0.05). For each site, the major and minor allele were inferred from genotype likelihoods. Allele frequencies were estimated both while assuming known major and minor alleles but also while taking the uncertainty of the minor allele inference into account (-doMajorMinor 1 -doMaf 3). Estimated variant calls for each chromosome were then used in a second step to produce a .bcf file with genotype likelihoods and posterior probabilities (-sites -doBcf 1 -doMajorMinor 3 -doMaf 1 -doPost 1 -GL 1) that was converted into .vcf with view in BCFtools (v.1.18)[7].

**6. SNP dataset for genomic diversity and GWAS**: Haplotype-phasing was performed in Beagle (v.4.1)[8] with default parameter settings, using genotype likelihoods to remove uncertainty in the initial .vcf file and add precision based on similarities between pairs of individuals. SNP imputation was performed in Beagle (v.5.2)[9] with default parameter settings, which are considered appropriate for a global population. Resulting per chromosome .vcf files were concatenated and indexed in BCFtools. To remove potential paralogs, sites with excess heterozygosity (flagged 'ExcHet<1') and with >5% heterozygote genotypes were filtered out. The resulting .vcf dataset contained 15,101,725 SNPs and was subsequently reformatted to .gds with the snpgdsVCF2GDS function (method = “biallelic.only”) in the R[10] package SNPRelate (v.0.9.19)[11].

**7. Environments of origin**: GPS coordinates of genotypes were taken directly from collection sites and used to extract data from raster files downloaded from the CHELSA v2.1 climate repository[12,13], and to create an elevation raster layer with the get_elev_raster function in the R package elevtr (v.0.99.0)[14]. Coordinates were then transformed to spatial points in the WGS84 Coordinate System (same as .tif raster files) with the SpatialPoints function in the R package sp (v.2.1-3)[15], and data from rasters was obtained using the extract function in the R package raster (v.3.6-26)[16]. The final environmental dataset included 52 variables (data S1) and environmental gradients across the cheatgrass distribution were identified with PCA (R function prcomp, variables scaled and centered) (fig. S6e). Furthermore, a shapefile of Level I Ecological Regions of North America[17] was downloaded from the USA Environmental Protection Agency. Data from this shapefile was extracted with the over function in the R package sp using the spatial points obtained above (data S1) to assign genotypes into ecological regions.

**8. SNP dataset for population genetic structure**: Using the dataset of 15,101,725 SNPs, sites in high linkage-disequilibrium were detected in PLINK (v.1.9)[18] with a window size of 150 kb, a step size of 1, and a pairwise $R^2$ threshold of 0.5. The pruned list of unlinked SNPs was then used to produce a .vcf with BCFtools view, containing 266,504 SNPs. This .vcf was then converted to .gds format (as in 6 above) for downstream analyses. To produce a genotype-likelihoods dataset, unlinked sites were used in ANGSD to estimate genotype likelihoods in the SAMtools framework, with the output written to Beagle (-GL 1 -doGlf 2 -doMajorMinor 1 -doMaf 3).

**9. Population structure analyses**: Cross-validation for the number of clusters was determined from log-likelihoods of the NGSadmix[19] output across all replicates[20]. For the PCA in PCAngsd (v.1.11)[21], individual allele frequencies were estimated on all sites in an iterative approach using a truncated singular value decomposition model, and the covariance matrix was estimated using the inferred individual allele frequencies from prior information for the unobserved genotypes. For plotting, samples were colored according to their admixture proportions using the function geom_scatterpie in the R package scatterpie (v.0.2.1), implemented in ggplot2[22]. To compute an unrooted phylogenetic tree with the Neighbor-Joining (NJ) algorithm[23], a genetic dissimilarity matrix was produced with the snpgdsDiss function in the R package SNPRelate and plotted with the plotnj function in the R package phyclust (v.0.1-33)[24] with tips colored according to their admixture proportions using the tiplabels function and the pies option in the R package ape (v.5.7-1)[25]. We calculated Nei's[26] pairwise $F_{ST}$ and Weir & Goudet's[27] population-specific $F_{ST}$ with functions pairwise.fst.dosage and fs.dosage in the R package hierfstat (v.0.5-11)[28]. For this, a genotype matrix with dosage data for all sites was estimated with the function snpgdsGetGeno in the R package SNPRelate.

**10. Variant annotation and genetic load**: We performed variant effect annotations with SnpEff (v 5.1f)[29]. First, we constructed a SnpEff database for *B. tectorum*. The reference genome and gene annotation files were downloaded from the Comparative Genomics platform CoGe (https://genomevolution.org/coge/), genome ID: id6435626. A coding sequence file was produced using gffread (v 0.12.8)[30] and the SnpEff annotation pipeline was applied to the .vcf of 307 genotypes and ~15M SNPs. Variants categorized as high-impact (chromosome large deletion, chromosome large duplication, chromosome large inversion, exon deleted, exon deleted partial, exon duplication, exon duplication partial, exon inversion, exon inversion partial, frame shift, gene deleted, gene fusion, gene fusion half, gene fusion reverse, gene rearrangement, protein-protein interaction locus, protein structural interaction locus, rare amino acid, splice site acceptor, splice, site donor, stop lost, stop gained, start lost, start gained, transcript deleted) or missense (non-synonymous) were subsequently identified and counted per genotype, along with synonymous variants.

**11. MEMs for spatial matrices**: Weighting matrices among unique sample locations were generated using the listw.candidates function in the adespatial (v.0.3-23) R package[31]. Two algorithms were implemented, Gabriel graph and distance-based graph, to generate three candidate connectivity matrices. The Gabriel graph results primarily in connections among neighboring sites. A distance-based graph connects sites closer than a given threshold, for which we used two values: minimum distance required to connect all points (*i.e.*, the largest distance of a minimum spanning tree) and infinity (resulting in a fully connected graph). With each of these three connectivity matrices, two spatial weighting matrices were generated using two distance-decay functions: Linear (weight between two sites = 1 − D/Dmax, where D is distance between sites and Dmax is maximum distance among all sites) or concave up (weight between two sites = D−0.01). Then, the forward selection of MEM eigenvectors algorithm was used to optimize the number of eigenvectors (restricted to those with positive eigenvalues) included in RDA for each MEM set[32]. Optimization is based on adjusted $R^2$, and the MEM set with greatest adjusted $R^2$ is defined as the optimal set. These eigenvectors were included in the RDA below on native and invaded whole-SNP datasets filtered for MAF>0.05 (native: 234,122 SNPs, invaded: 212,292 SNPs).

**12. Phenotypes**: During the grow out in 2020, we measured phenotypes on up to 184 genotypes with 2–3 replicates that emerged within ~9–18 days of planting and survived until harvesting. Plants were monitored every two days until phenotyping was terminated (i.e., termination: ~250 days after germination when >90% of plants had flowered), and then once a week until the last plants were harvested. Eleven phenotypes were recorded: seedling and adult (i.e., reproductive) height (used to get spring growth), number of leaves, number of tillers, days to flower, inflorescence height, dry biomass, total seed mass (i.e., fecundity), individual seed mass (i.e., seed mass), total seed length, and awn length. No phenotypes were recorded after termination, but seed data were recorded for 11 extra genotypes that had not flowered prior to termination.

Seedling and adult height were measured in centimeters from the base of the plant to the tip of the longest leaf. Seedling height was measured the day plants were taken out of vernalization, *i.e.* 12 weeks after planting. Adult height was taken from reproductive plants, close to or at harvesting. Spring growth (cm) was the growth of plants after vernalization, taken as the difference between adult and seedling height. Number of leaves and tillers were counted when adult height was measured in reproductive plants. Days to flower were counted from the day of individual germination to the day awn tips appeared through the boot and was recorded until termination. Harvesting was performed when >75% of the main inflorescence was matured; at this point inflorescence height was measured in centimeters from the base of the plants to the tip of the tallest panicle. Uncleaned seeds were collected, stored in coin envelopes, kept under dry conditions for ~30 days, and weighed in milligrams in a tared balance. Vegetative tissue was cut at the soil surface, collected in paper bags, oven-dried for 48 hours at 38ºC, and weighed in milligrams in a tared balance. The sum of these two weights constituted total dry biomass. Fecundity and seed mass were determined for a single replicate per genotype due to the time intensity involved in this task. Viable seeds (*i.e.*, filled) were cleaned and weighed in milligrams in a tared balance, constituting fecundity, then 20 random seeds were weighted to get an average seed mass. Seed and awn length were measured with a caliper in five seeds per replicate. Seed length was measured from the base of the raquilla to the tip of the awn, and awn length was measured from the tip of the palea to the tip of the awn.

As mentioned in *Plant material*, for 155 genotypes planted seeds were directly collected from the field, thus maternal environment effects could be a possible source of phenotypic variation. However, we found a strong correlation between growth chamber and common garden flowering time (common garden planted in 2021 from S1 seeds). Flowering time was averaged at the genotype level from the raw data for each study (*i.e.*, growth chamber and common garden). The Pearson and Spearman correlation coefficients were 0.65 and 0.58, respectively, suggesting little maternal effects in growth chamber data.

**13. Common Gardens**: For each growing season, we planted seeds directly on the ground in the fall of the year preceding the growing season. To track individual plants, seeds were glued to a toothpick (detailed protocol in Vahsen et al. 2025[33]). In the 2022 growing season, we planted 100 plants each per plot in a randomized blocked design, with each density and temperature treatment combination represented once across 10 total blocks per site, for a total of 8,000 plants (2 density treatments × 2 gravel treatments × 10 blocks × 100 plants × 2 common garden sites = 8,000 total plants[33]). In the 2023 growing season, we reduced the replication at the plot level, such that at the cold, less seasonal site, there were 80 plants per plot (3,200 total plants) and at the warm, seasonal site, there were 90 plants per plot (3,600 total plants). Plants were not irrigated throughout the experiment. We recorded emergence starting in early winter the year preceding the growing season and recorded individual plant growing stage on a roughly biweekly basis starting in the spring of the growing season. We considered plants to be flowering for this analysis when their florets were first observed to have emerged and were green in color.

At the end of the growing season, we opportunistically harvested the aboveground biomass of plants so that florets had a chance to mature (shift from green to purple in color), while avoiding plant senescence and the dropping of seeds. Plant harvesting occurred within the following date ranges for each common garden site and year: cold, less seasonal site = June 23 – July 8, 2022 and June 14 – August 4, 2023; warm, seasonal site = February 24 – June 28, 2022 and March 4 – June 11, 2023. We stored the aboveground biomass of each individual plant in a separate paper envelope at room temperature in the lab prior to processing seed mass and weight.

For processing in 2022, we hand-separated each plant into vegetative and reproductive biomass. Depending on the total number of seeds, we then either separated, counted, and weighed all individual seeds (i.e., if $n < 50$), or we took a subset of 50 seeds from the total number of seeds and recorded their weight. Then, for those plants for which only a subset of seeds was counted and weighed, we estimated the total seed count given the weight of the seed subset and the total reproductive biomass. For processing in 2023, we only hand-separated each plant into vegetative and reproductive biomass and did not count the seeds. Thus, the fitness metric reported for the 2022 data is total seed count and the fitness metric reported for the 2023 data is reproductive biomass. Plants that were infested with smut or were noted to have dropped seeds prior to or during harvest were not included in the analyses.

**14. Inter-chromosomal linkage-disequilibrium (LD)**: Because mean LD was high even at 5Mb, we also assessed inter-chromosomal LD. To this end, we randomly sampled ~15K genome-wide SNPs and used the program GWLD[34] to get an $R^2$ between all pairs of SNPs (with MAF>0.05) in different chromosomes. Using the *Bromus tectorum* CoGe annotations, top GWAS SNPs (those with the lowest 100 *p*-values for each GWAS) were assigned to candidate genes using a 200 kb window based on the observed LD decay. Given the range-wide LD we observed, we also used a 1Mb window, but no other clear QTL were revealed. We thus report QTL using a 200 kb window.

### Supplementary Table

**Supplementary Table 1. Variation of growth chamber phenotypes.** Broad-sense heritability ($H^2$) and variation of best-linear-unbiased-estimates (BLUE) in eleven growth chamber phenotypes.

| Trait | $H^2$ | $BLUE_{mean}$ | $BLUE_{sd}$ | $BLUE_{min}$ | $BLUE_{max}$ |
|---|---|---|---|---|---|
| Tillers number | 0.54 | 6.85 | 1.76 | 2.82 | 11.22 |
| Leaves number | 0.70 | 42.96 | 19.38 | 0 | 92.39 |
| Adult height (cm) | 0.43 | 16.38 | 2.41 | 8.97 | 23.70 |
| Spring growth (cm) | 0.45 | 9.21 | 2.84 | 0.85 | 17.68 |
| Biomass (mg) | 0.45 | 707.64 | 230.56 | 218.01 | 1321.59 |
| Days to flower | 0.92 | 174.76 | 33.31 | 104.07 | 250.58 |
| Inflorescence height (cm) | 0.68 | 34.11 | 9.38 | 11.18 | 60.21 |
| Fecundity (mg) | 0.55 | 282.56 | 165.21 | 6.02 | 1003.02 |
| Seed mass (mg) | 0.92 | 2.67 | 0.73 | 1.11 | 6.18 |
| Seed length (mm) | 0.72 | 26.77 | 3.12 | 20.16 | 36.07 |
| Awn length (mm) | 0.66 | 17.46 | 2.64 | 11.57 | 23.16 |

**Supplementary Figures**

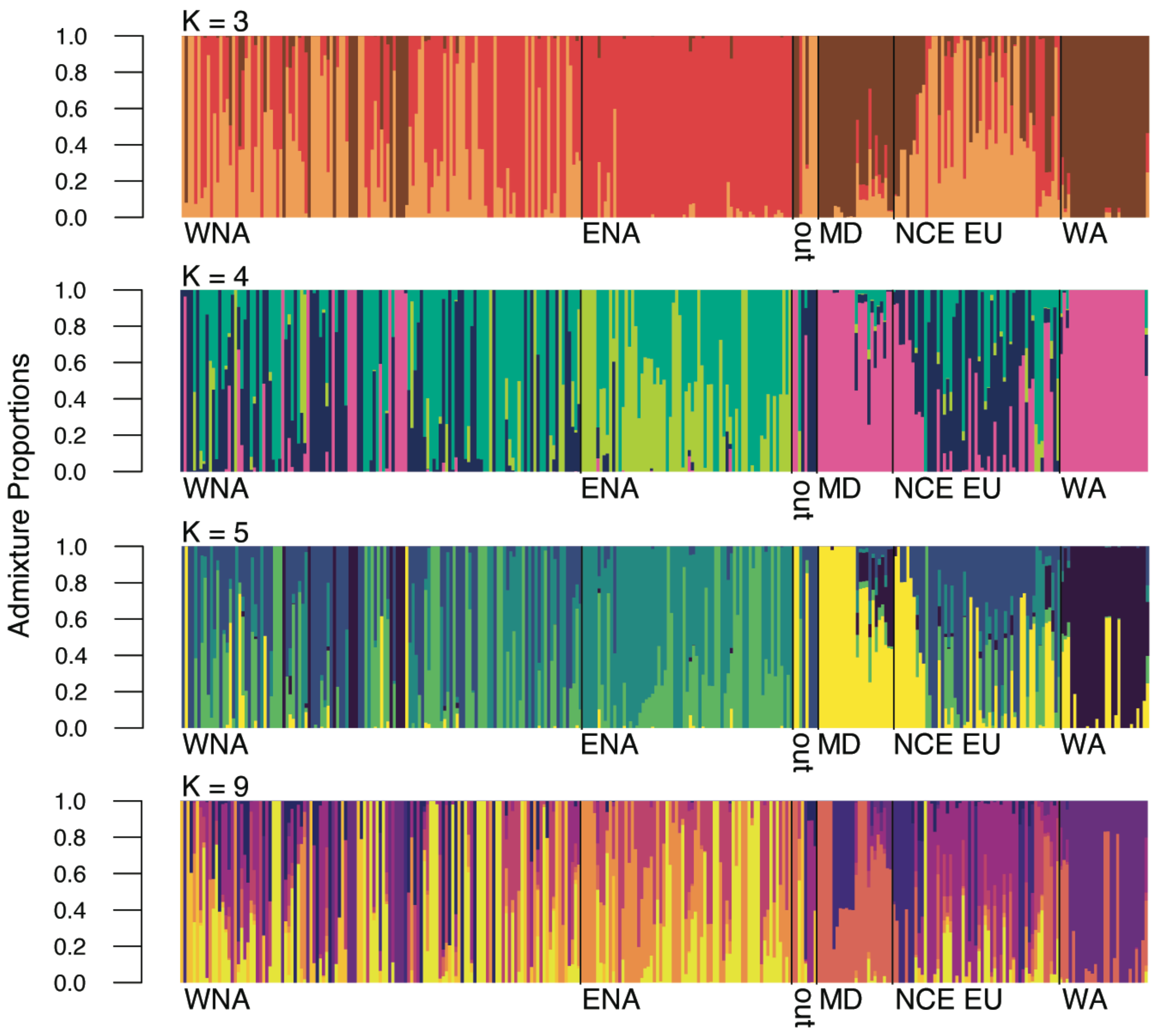


**Supplementary Fig. 1. NGSadmix results for K=3, K=4, K=5, and K=9 ancestral genetic clusters.** Bars along the x-axis are the 307 studied genotypes arranged by longitude and separated by region of origin. Invasive: WNA=western North America (n=107), ENA=eastern North America (n=67), out=outside of North America (n=8). Native: MD=Mediterranean (n=24), NCE EU= north-central-east Europe (n=53), WA=west Asia (n=28).

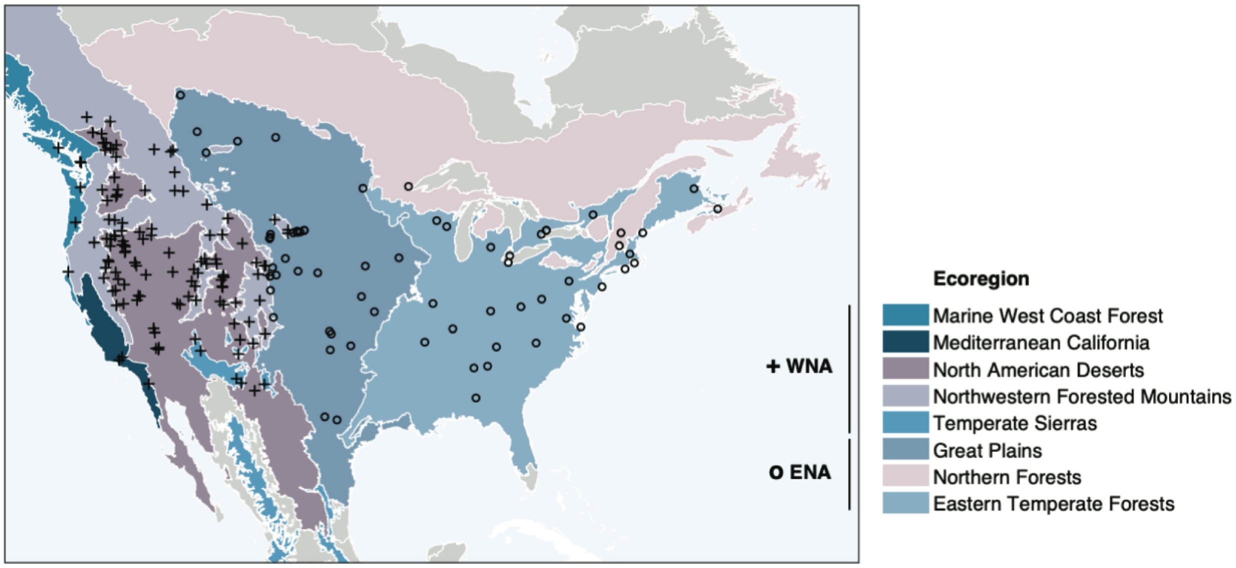


**Supplementary Fig. 2. Invasive genotypes from North America with ecoregion of origin.** Map of western North America (WNA) and eastern North America (ENA) invaded range genotypes, with colors showing the Level I Ecoregions of North America[17] classification used to delimit WNA and ENA.

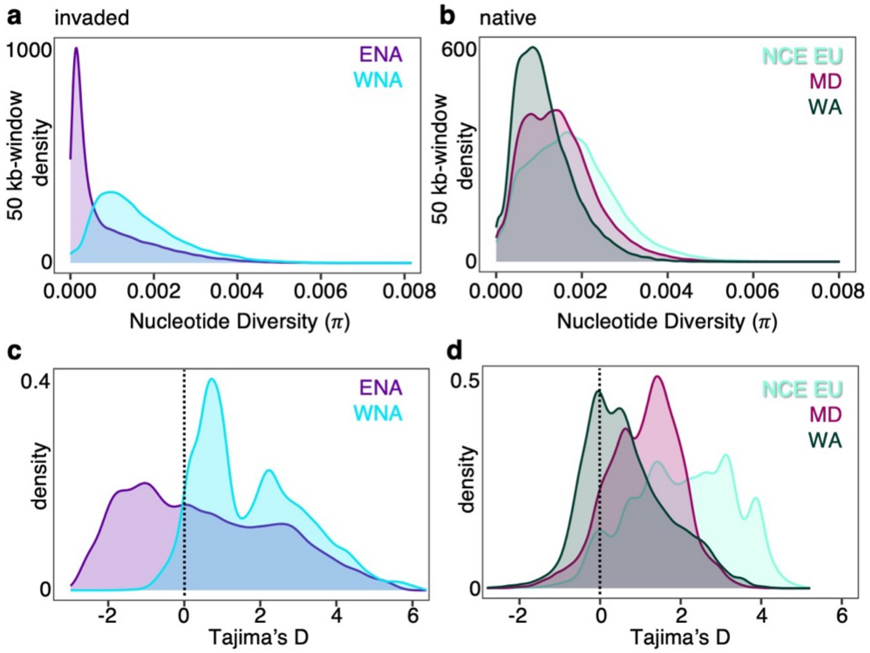


**Supplementary Fig. 3. Patterns of genomic diversity in genotypes from different regions.** (**a** and **b**) Nucleotide diversity (π) based on ~15M sites. (**c** and **d**) Tajima's D based on shared polymorphisms across ~15M sites. Colors in each panel correspond to regions in the invaded (WNA=western North America, ENA=eastern North America) and the native range (NCE EU= north-central-east Europe, MD=Mediterranean, WA=west Asia). Genome-wide estimates were obtained from 50 kb sampling windows.

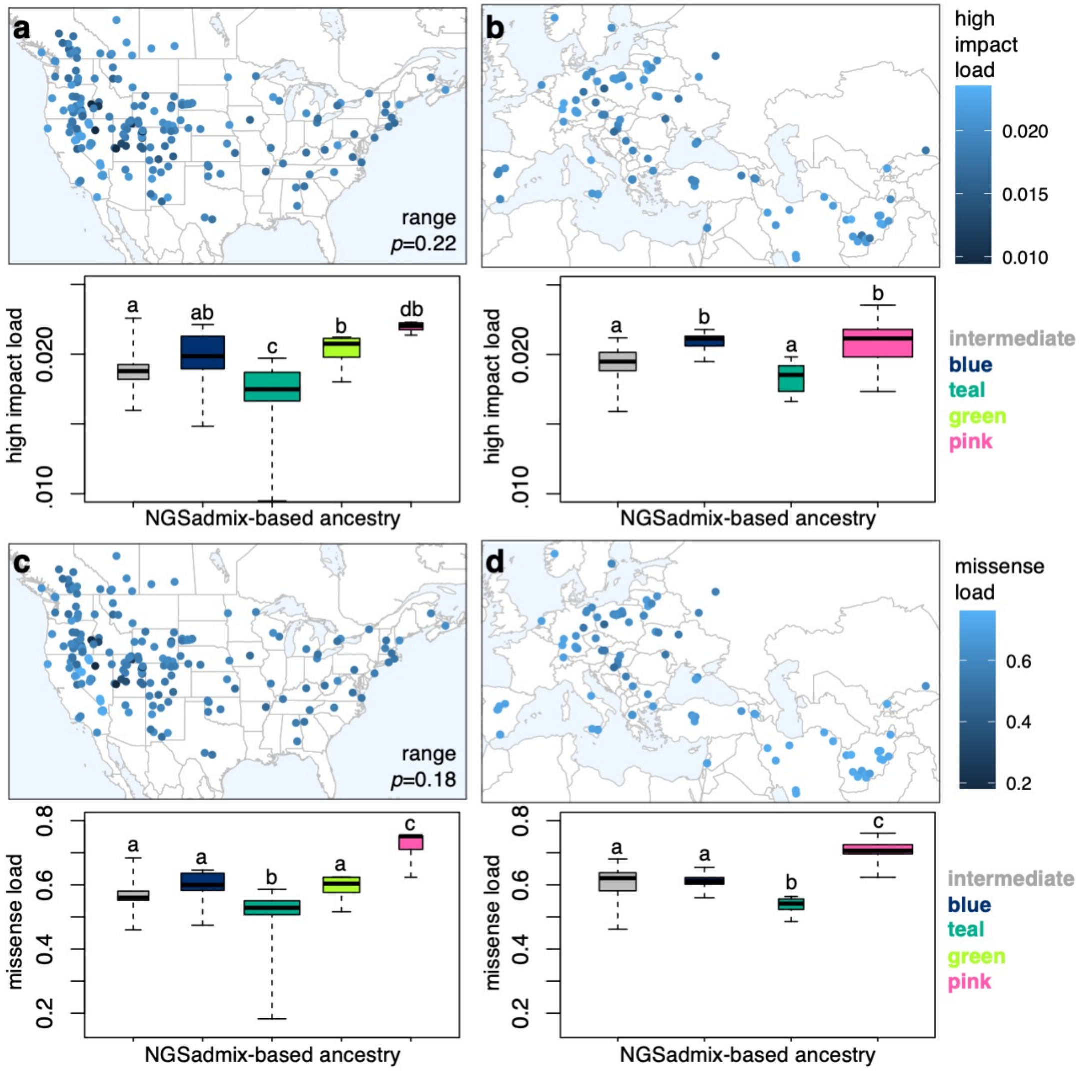
a
b
c
d
range
p=0.22
range
p=0.18
high impact load
0.020
0.015
0.010
missense load
0.6
0.4
0.2
high impact load
0.020
.010
a
ab
c
b
db
a
b
a
b
NGSadmix-based ancestry
intermediate
blue
teal
green
pink
missense load
0.8
0.6
0.4
0.2
a
a
b
a
c
a
a
b
c
NGSadmix-based ancestry
intermediate
blue
teal
green
pink

**Supplementary Fig. 4. Range-wide patterns of estimated genetic load.** Geographic distribution and variation of normalized high impact load in (**a**) invasive and (**b**) native genotypes. Geographic distribution and variation of normalized missense (*i.e.*, non-synonymous) load in (**c**) invasive and (**d**) native genotypes. Boxplots are colored based on K=4 ancestral genetic clusters as in Fig. 1 (with names used in the main text to the right). Genotypes were assigned to a cluster when they had >0.55 NGSadmix ancestry proportion, if neither ancestry was >0.55 then they were assigned as intermediate. Boxplots width is proportional to sample size. Boxplots indicate median (middle line), $25^{th}$, $75^{th}$ percentile (box), and whiskers cover the data extent. Range differences (noted in maps to the left) and ancestry group differences *p*-values come from a Tukey HSD test on a 2-way ANOVA that found no significant differences ($p$=0.3–1) between native and invaded genotypes of the same ancestry. A shared letter (or letter combination) indicates no significant differences ($p$>0.05) between ancestry groups. The light green ancestry is only present in the invaded range because no genotypes had >0.55 of it in the native range.

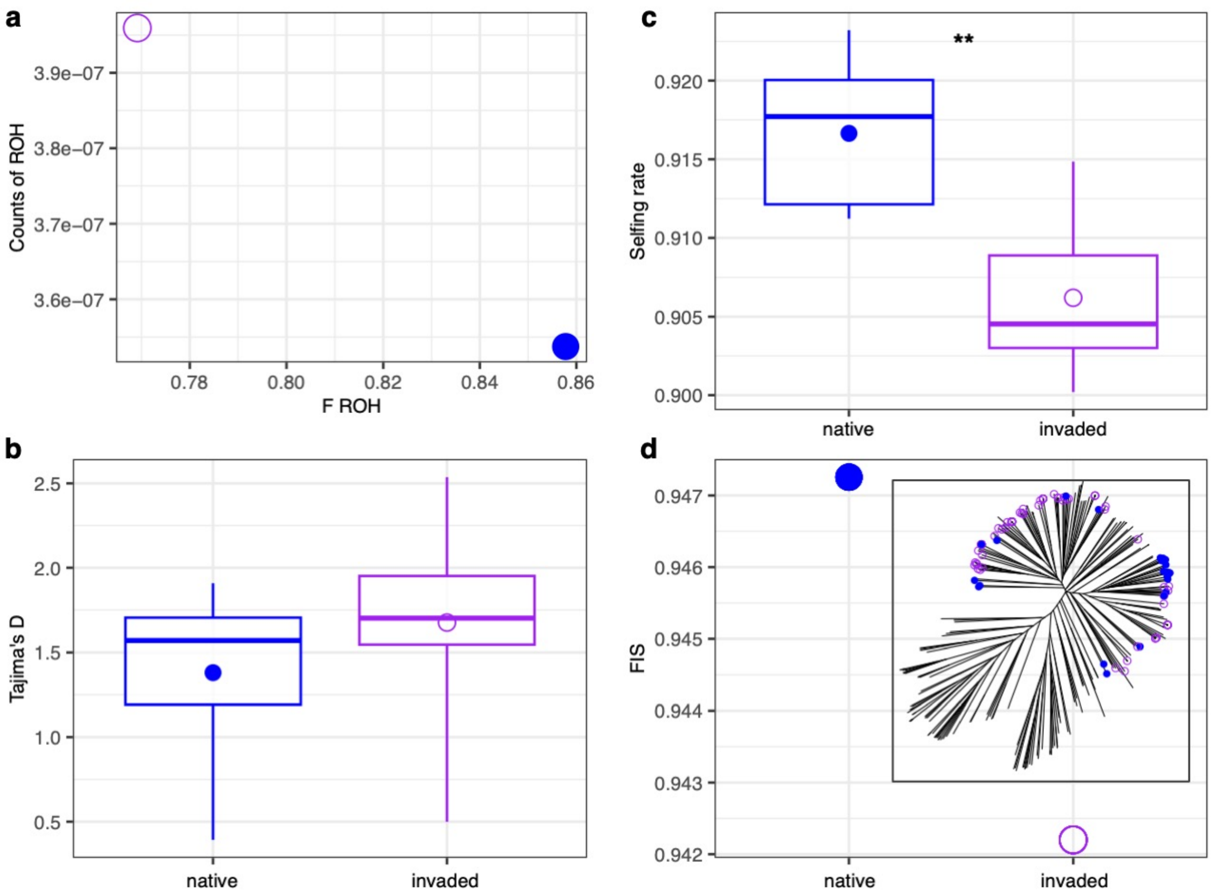


**Supplementary Fig. 5. No evidence of increased inbreeding due to selfing in the invaded relative to the native range**. (**a**) The pattern of runs of homozygosity (ROH) represented by the counts of ROH relative to the proportion of the genome in ROH (F ROH) in native (closed circle) and invaded (open circle) range genotypes from the same lineage. Values are means across the seven cheatgrass chromosomes for each group using ~15M genome-wide sites. Estimated: (**b**) Tajima's D, (**c**) Selfing rates (***$p$=0.002), and (**d**) Inbreeding coefficients ($F_{IS}$) for the same groups using the same loci. Boxplots represent variation across the seven cheatgrass chromosomes and indicate median (middle line), 25th, 75th percentile (box), and whiskers cover the data extent. Circles are means. The inset neighbor-joining tree in (d) denotes the lineage chosen to compare closely related native (n=27) and invasive (n=74) genotypes that were sequenced from seedlings of seeds removed directly from the field.

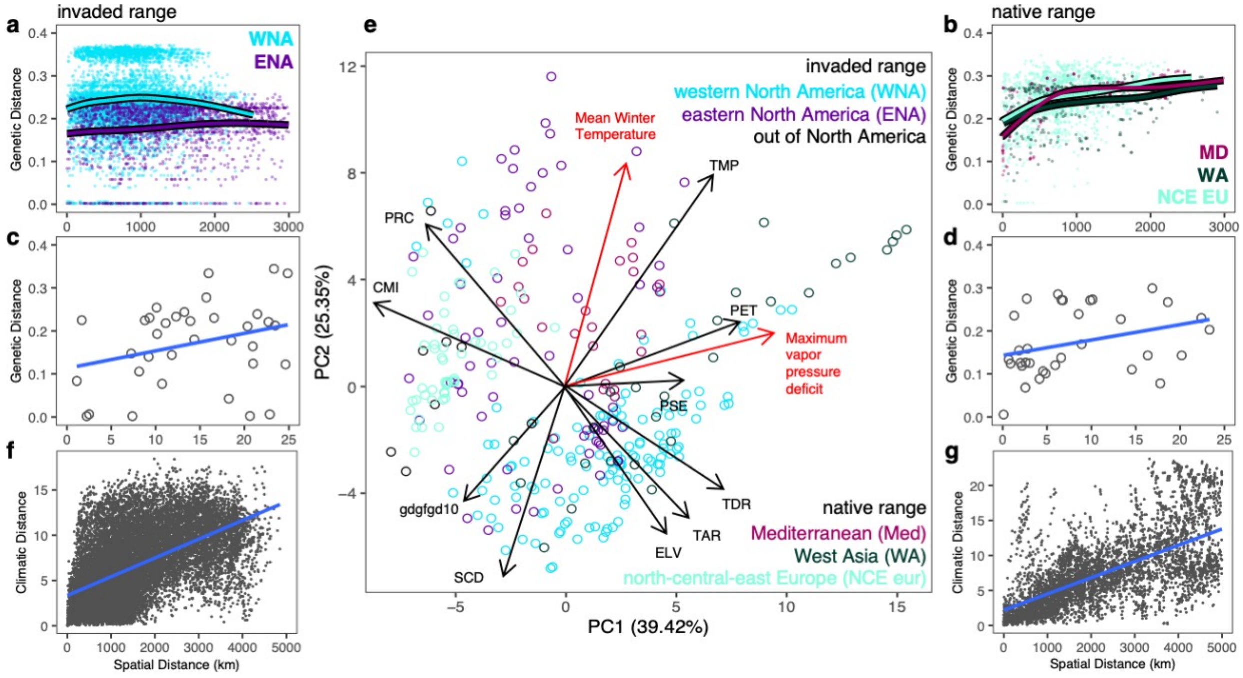


**Supplementary Fig. 6. Genetic and climatic spatial variation in study genotypes.** (**a**) Great spatial genetic heterogeneity at short geographic distance in western North America (WNA), but not in eastern North America (ENA) and (**b**) significant isolation-by-distance (IBD) in all native range regions. (**c**) At small scales (0–25 km), IBD remains weak in the invaded range, and (**d**) strong in the native range. (**e**) Eigenvector plot of the loadings of 52 abiotic CHELSA variables onto PC1 and PC2 describing environmental variation in 307 genotypes colored by region. Eigenvectors of selected variables are depicted based on their large loadings into each PC, with variables in red having the largest loading. CMI: mean monthly climate moisture index, ELV: elevation, gdgfgd10: first growing degree day above 10. PET: potential evapotranspiration, PRC: total annual precipitation, PSE: precipitation seasonality, SCD: snow cover days, TAR: temperature annual range, TDR: temperature diurnal range, TMP: annual mean temperature. (**f**) Great spatial environmental heterogeneity at short distance in the invaded range. (**g**) Environmental variation generally increases with distance in the native range.

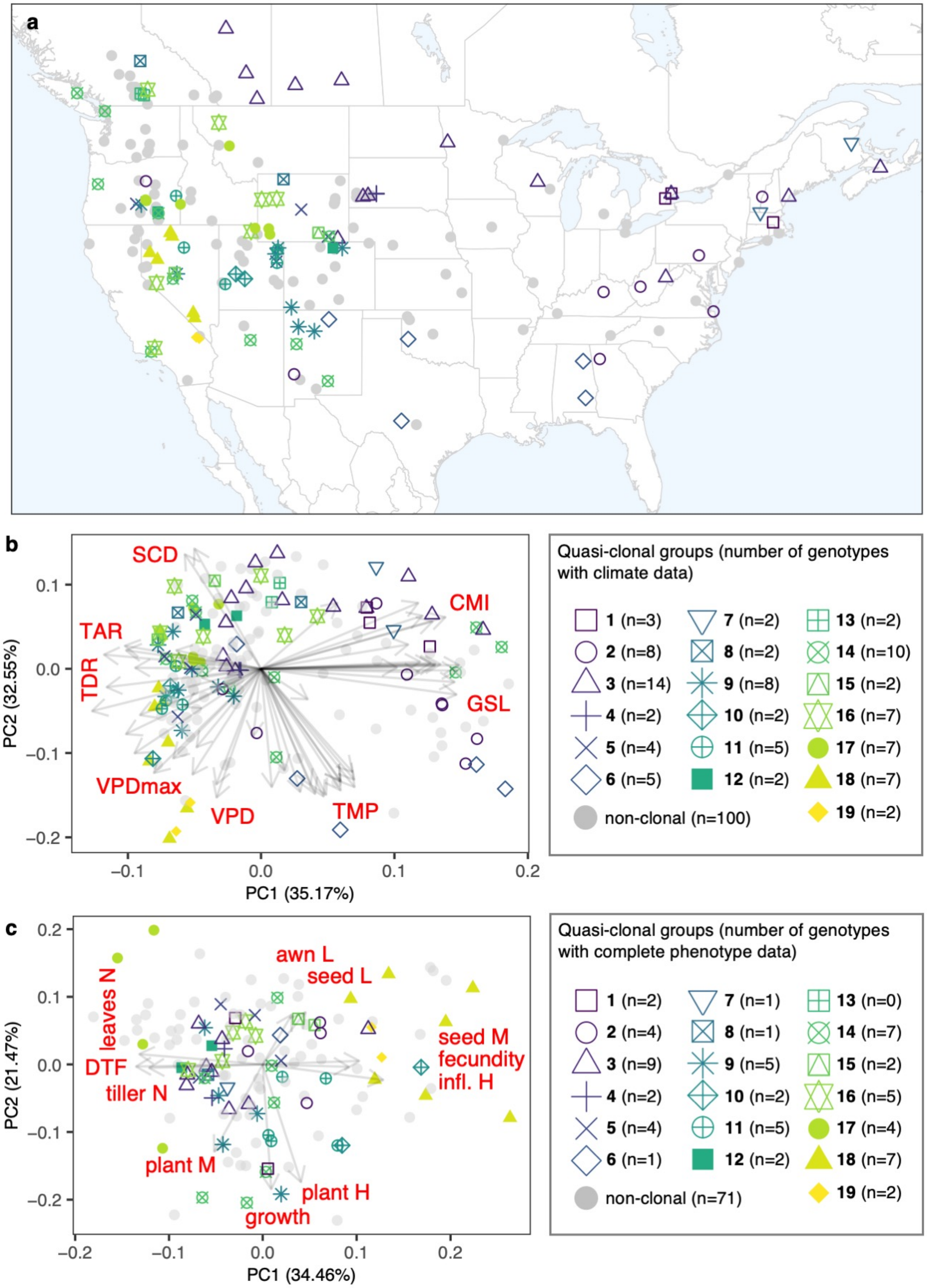

a
b
SCD
TAR
TDR
CMI
GSL
VPDmax
VPD
TMP
PC2 (32.55%)
PC1 (35.17%)
0.1
0.0
−0.1
−0.2
0.2
Quasi-clonal groups (number of genotypes with climate data)
1 (n=3)
2 (n=8)
3 (n=14)
4 (n=2)
5 (n=4)
6 (n=5)
7 (n=2)
8 (n=2)
9 (n=8)
10 (n=2)
11 (n=5)
12 (n=2)
13 (n=2)
14 (n=10)
15 (n=2)
16 (n=7)
17 (n=7)
18 (n=7)
19 (n=2)
non-clonal (n=100)
c
awn L
seed L
leaves N
DTF
tiller N
seed M
fecundity
infl. H
plant M
plant H
growth
PC2 (21.47%)
PC1 (34.46%)
Quasi-clonal groups (number of genotypes with complete phenotype data)
1 (n=2)
2 (n=4)
3 (n=9)
4 (n=2)
5 (n=4)
6 (n=1)
7 (n=1)
8 (n=1)
9 (n=5)
10 (n=2)
11 (n=5)
12 (n=2)
13 (n=0)
14 (n=7)
15 (n=2)
16 (n=5)
17 (n=4)
18 (n=7)
19 (n=2)
non-clonal (n=71)

**Supplementary Fig. 7. Genetically distinct groups of near-clonal genotypes can be geographically widespread but remain climatically and phenotypically differentiated.** (**a**) Map of the 194 North American genotypes, marked by group assignment following legend in b. (**b**) Eigenvector plot of the loadings of 52 abiotic CHELSA[12,13] variables onto PC1 and PC2 describing environmental variation in the 194 North American genotypes**.** Eigenvectors are depicted in light grey, with variables in red having the largest loadings onto PC1 or PC2. CMI: mean monthly climate moisture index, GSL: growing season length, SCD: snow cover days, TAR: temperature annual range, TDR: temperature diurnal range, TMP: annual mean temperature, VPD: vapor pressure deficit, VPDmax: maximum vapor pressure deficit. (**c**) Eigenvector plot of the loadings of 11 growth chamber phenotypes describing trait variation in 135 North American genotypes. Eigenvectors are depicted in light grey for variables; awn L: awn length, seed L: seed length, seed M: individual seed mass, fecundity: total seed mass, infl. H: inflorescence height, plant H: plant height, growth: growth in height after vernalization (spring growth), plant M: total dry aboveground biomass, tiller N: number of tillers, DTF: days to flower, leaves N: number of leaves.

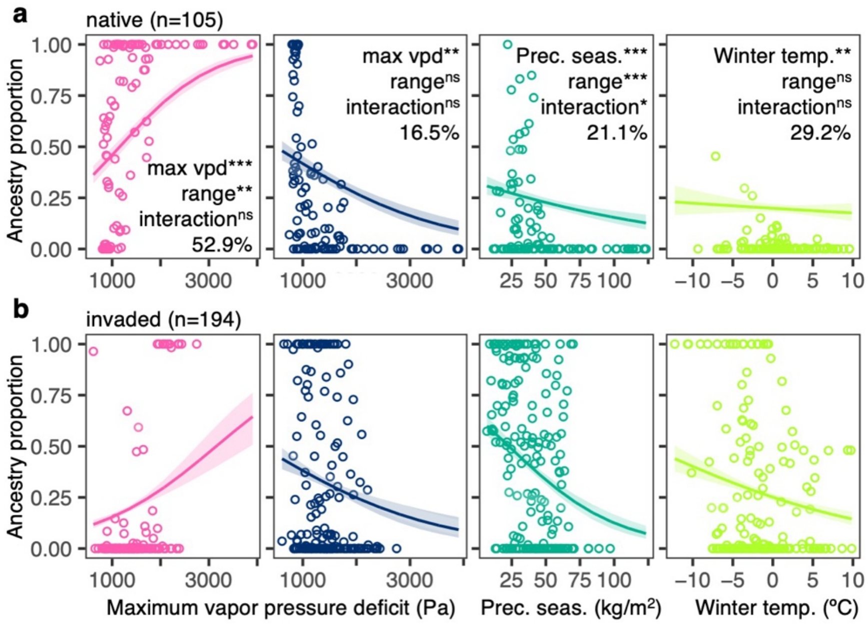


**Supplementary Fig. 8. Ancestry-environment associations in the native range are mirrored in North America.** Ancestry-environment associations in (**a**) native and (**b**) invaded range genotypes, showing results of generalized-additive-models (GAM) where predictors were an environmental gradient, range, and their interaction: $^{ns}p>0.05$, $^{**}p<0.005$, $^{***}p<0.0005$; % denotes model deviance. Colors are K=4 ancestral clusters. max vpd: maximum vapor pressure deficit, Prec. seas.: precipitation seasonality, temp.: mean temperature.

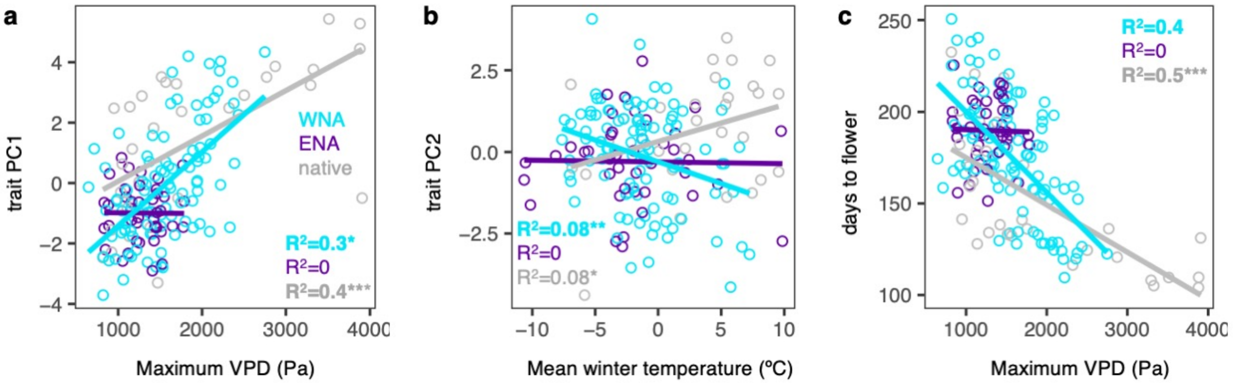


**Supplementary Fig. 9. Trait-environment clines are absent in ENA.**

(**a** to **c**) Growth chamber phenotype-environment associations for genotypes in western North America (WNA, n=95), eastern North America (ENA, n=41), and the native range (n=31), examined in Fig. 3B. Coefficients of determination ($R^2$) and trends (lines) come from linear regressions with $R^2$ highlighted if significant ($p$<0.05). Asterisks represent significance from linear-mixed kinship models that accounted for relatedness among genotypes: *$p$<0.05, **$p$<0.005, ***$p$<0.0001.

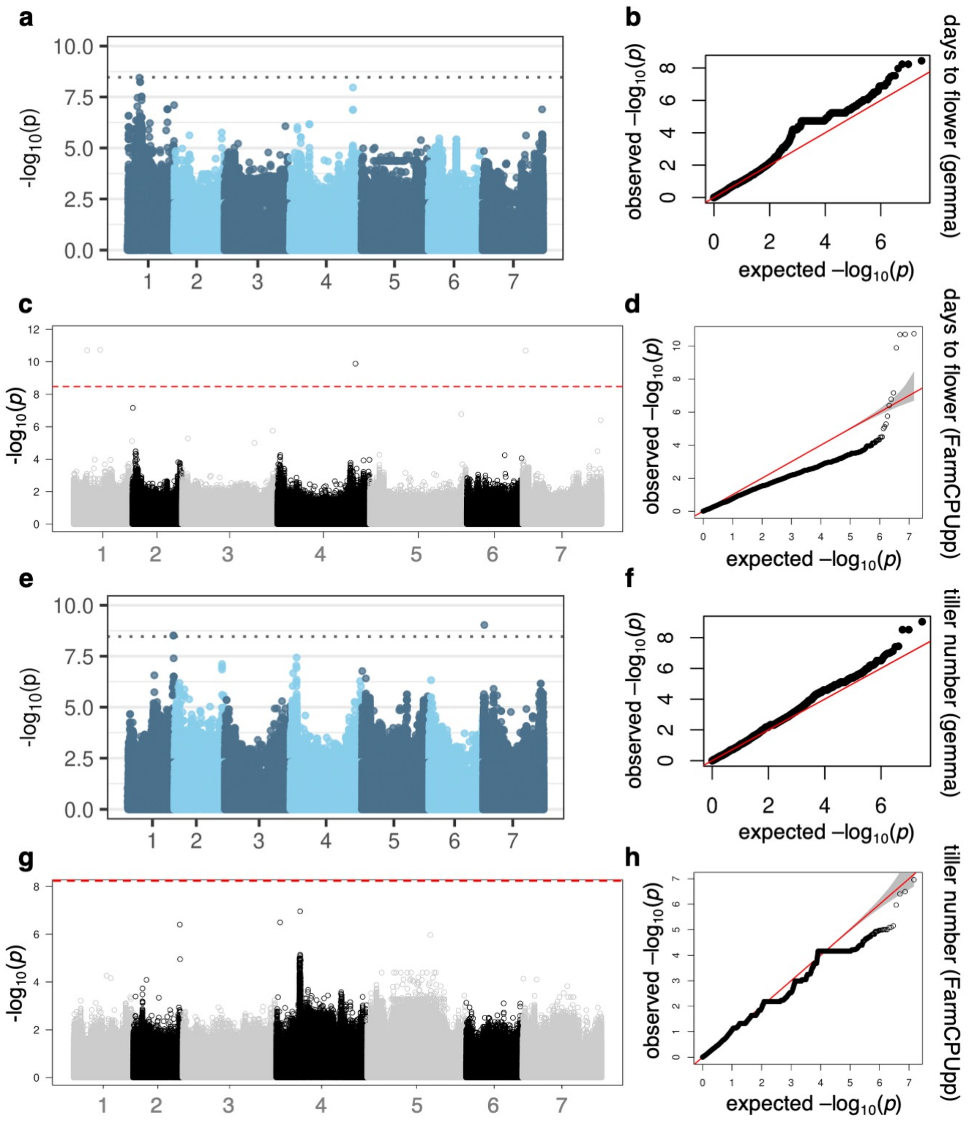


**Supplementary Fig. 10. Results of genome-wide association studies (GWAS) in two growth chamber phenotypes using two methods.** Manhattan plots with Bonferroni line (left) and respective quantile-quantile plot (right). Phenotype and method are shown to the right. Flowering time GWAS with (**a**, **b**) univariate linear mixed model (LMM, gemma) and (**c**, **d**) multi-locus mixed model (MLMM, FarmCPUpp) performed on 173 genotypes and 14,710,773 SNPs filtered for MAF<0.05. Tiller number GWAS with (**e**, **f**) LMM and (**g**, **h**) MLMM performed on 184 genotypes and 14,632,788 SNPs filtered for MAF<0.05.

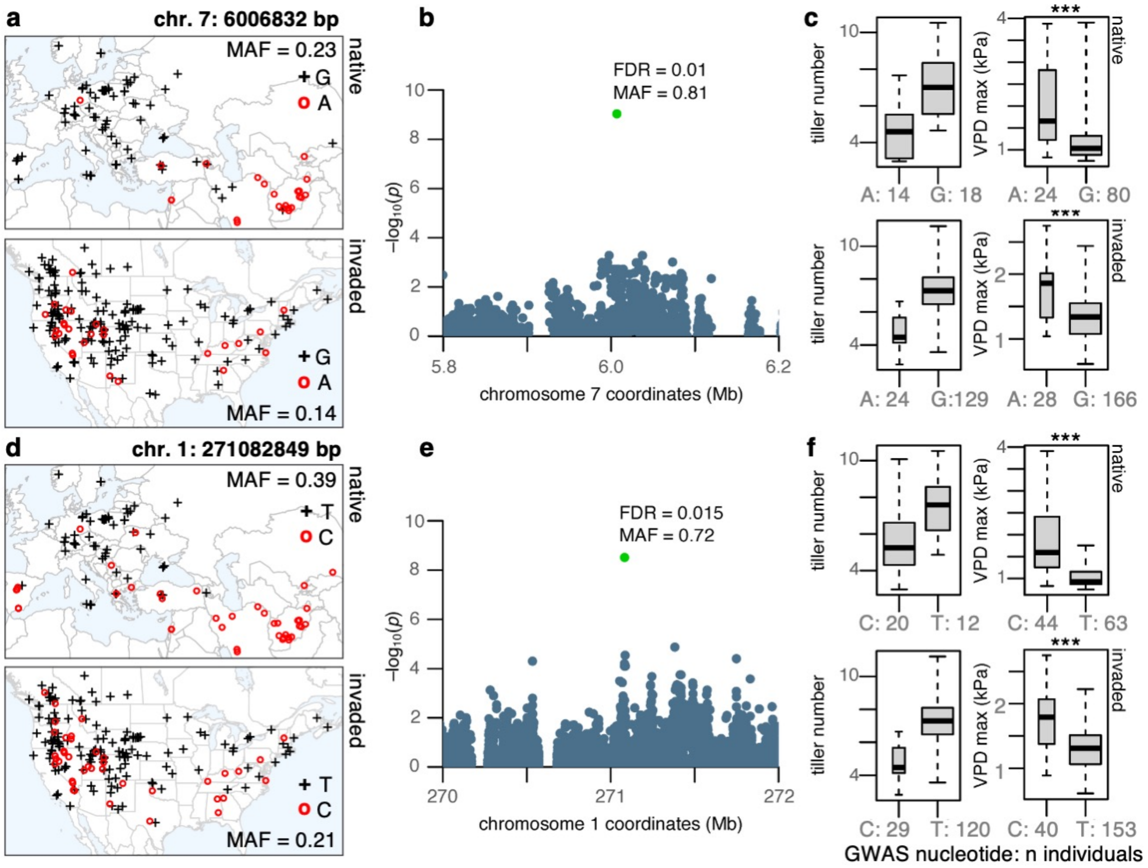


**Supplementary Fig. 11. Environmental trends of two tillering QTL are mirrored between continents.** (**a** and **d**) Geographic distribution of QTL SNP alleles in the native (top) and invaded (bottom) range; crosses represent the reference/major allele, and open circles the alternate/minor allele. (**b** and **e**) Zoomed-in Manhattan plots showing Wald-test *p*-values (plotted as $-\log_{10}$) from GWAS and genomic location of top SNP (marked in green), with respective false-discovery-rate (FDR) and minor allele frequency (MAF). (**c** and **f**) Phenotypic (boxplots to the left) and environmental variation (boxplots to the right) of flowering time QTL SNP alleles identified with GWAS. Boxplots indicate median (middle line), $25^{th}$, $75^{th}$ percentile (box), and whiskers cover the data extent. ***$p<0.001$ from two-tailed t-tests, but kinship linear-mixed models showed no significant differences. Max VPD: Maximum vapor pressure deficit in kPa. G, A, T, C on maps and x-axis of boxplots indicate nucleotides.

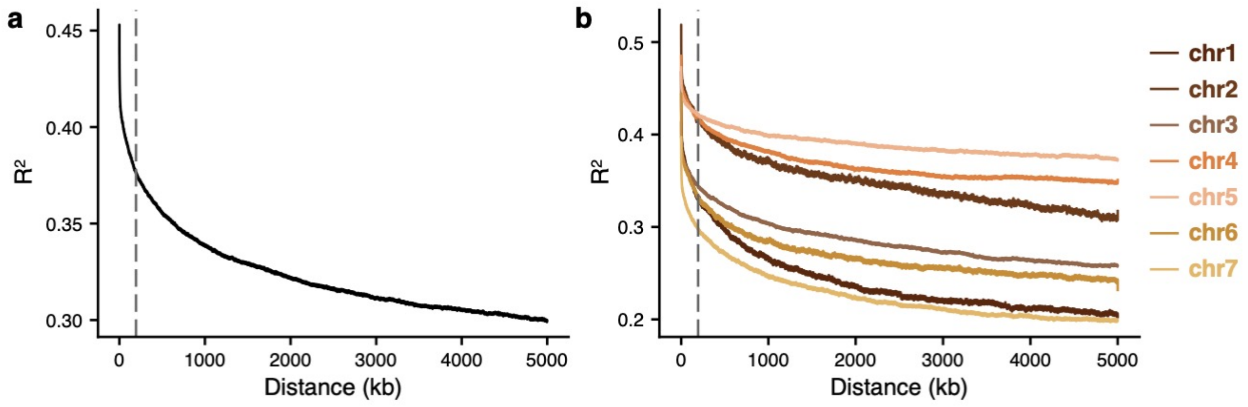


**Supplementary Fig. 12. Linkage disequilibrium (LD) as a function of genomic distance in *Bromus tectorum***. Using 307 genotypes and ~15M genome-wide SNPs filtered for MAF<0.05, LD was computed (**a**) genome-wide and (**b**) separately for all SNPs in each of the seven chromosomes. LD was computed as the square of the Pearson correlation between all pairs of SNPs ($R^2$) separated by up to 5 Mb. LD decay is plotted as the mean LD in genomic bins of 10 bp (from 0 – 100 bp) and of 100 bp (from 100 bp – 5 Mb). The dashed vertical lines indicate the distance at which the maximum mean LD decayed to ½ of the mean LD observed at 5 Mb.
